# Longitudinal isolation of potent near-germline SARS-CoV-2-neutralizing antibodies from COVID-19 patients

**DOI:** 10.1101/2020.06.12.146290

**Authors:** Christoph Kreer, Matthias Zehner, Timm Weber, Cornelius Rohde, Sandro Halwe, Meryem S. Ercanoglu, Lutz Gieselmann, Michael Korenkov, Henning Gruell, Philipp Schommers, Kanika Vanshylla, Veronica Di Cristanziano, Hanna Janicki, Reinhild Brinker, Artem Ashurov, Verena Krähling, Alexandra Kupke, Hadas Cohen-Dvashi, Manuel Koch, Simone Lederer, Nico Pfeifer, Timo Wolf, Maria J.G.T. Vehreschild, Clemens Wendtner, Ron Diskin, Stephan Becker, Florian Klein

**Author notes:** These authors contributed equally (Christoph Kreer, Matthias Zehner).

## Abstract

The SARS-CoV-2 pandemic has unprecedented implications for public health, social life, and world economy. Since approved drugs and vaccines are not available, new options for COVID-19 treatment and prevention are highly demanded. To identify SARS-CoV-2 neutralizing antibodies, we analysed the antibody response of 12 COVID-19 patients from 8 to 69 days post diagnosis. By screening 4,313 SARS-CoV-2-reactive B cells, we isolated 255 antibodies from different time points as early as 8 days post diagnosis. Among these, 28 potently neutralized authentic SARS-CoV-2 (IC_100_ as low as 0.04 μg/ml), showing a broad spectrum of V genes and low levels of somatic mutations. Interestingly, potential precursors were identified in naïve B cell repertoires from 48 healthy individuals that were sampled before the COVID-19 pandemic. Our results demonstrate that SARS-CoV-2 neutralizing antibodies are readily generated from a diverse pool of precursors, fostering the hope of rapid induction of a protective immune response upon vaccination.

## INTRODUCTION

By mid-May 2020 over 4.5 million severe acute respiratory syndrome coronavirus 2 (SARS-CoV-2) infections and over 300,000 casualties of the associated coronavirus disease 2019 (COVID-19) were reported (Dong et al., 2020; Huang et al., 2020; Zhou et al., 2020; Zhu et al., 2020). The exponential spread of the virus has caused countries to shut down public life with unprecedented social and economic consequences. Therefore, decoding SARS-CoV-2 immunity to promote the development of vaccines as well as potent antiviral drugs is an urgent health need (Sanders et al., 2020).

Monoclonal antibodies (mAbs) have been demonstrated to effectively target and neutralize viruses such as Ebola virus (EBOV; Ehrhardt et al., 2019; Flyak et al., 2016; Saphire et al., 2018), respiratory syncytial virus (RSV; Kwakkenbos et al., 2010), influenza virus (Corti et al., 2011; Joyce et al., 2016; Kallewaard et al., 2016), or human immunodeficiency virus 1 (HIV-1; Burton et al., 2009; Huang et al., 2016a, 2016b; Scheid et al., 2011; Schommers et al., 2020; Wu et al., 2010). The most prominent target for an antibody-mediated response on the surface of SARS-CoV-2 virions is the homotrimeric spike (S) protein. The S protein promotes cell entry through the interaction of a receptor-binding domain (RBD) with angiotensin-converting enzyme 2 (ACE2; Hoffmann et al., 2020; Walls et al., 2020). Antibodies that target the S protein are therefore of high value to prevent and treat COVID-19 (Burton and Walker, 2020).

## RESULTS

### SARS-CoV-2-infected individuals develop a polyclonal memory B cell response against the S protein

To investigate the antibody response against SARS-CoV-2, we collected blood samples from seven COVID-19 patients (aged 38 to 59 years) between 8 and 36 days post diagnosis (**Figure 1A, Table S1**). Five patients presented with mild symptoms including dry cough, fever, and dyspnoea, while two patients were asymptomatic (**Table S1**). Purified plasma immunoglobulin G (IgG) of all seven individuals showed binding to the full trimeric S-ectodomain (Wrapp et al., 2020), with half maximal effective concentrations (EC_50_) ranging from 3.1 to 96.1 μg/ml (**Figure 1B, Table S2**). Moreover, neutralizing IgG activity was determined against authentic SARS-CoV-2, showing 100% inhibitory concentrations (IC_100_) between 78.8 and 1,500 μg/ml in five out of seven patients (**Figure 1B, Table S2**). In order to decipher the SARS-CoV-2 B cell and antibody response on a molecular level, we performed single B cell sorting and sequence analysis of all individuals. Using flow cytometry, we detected between 0.04% (± 0.06) and 1.02% (± 0.11) IgG^+^ B cells that reacted with the S-ectodomain (**Figure 1C, Figure S1**). From these we isolated a total of 1,751 single B cells and amplified IgG heavy and light chains using optimized PCR protocols (**Figure 1C, Table S3**; Kreer et al., 2020a; Schommers et al., 2020). Sequence analysis revealed a polyclonal antibody response with 22% to 45% clonally related sequences per individual and 2 to 29 members per identified B cell clone (**Figure 1D, Table S3**). We conclude that a polyclonal B cell response against the SARS-CoV-2 S protein was initiated in all studied COVID-19 patients.

**Figure 1.**
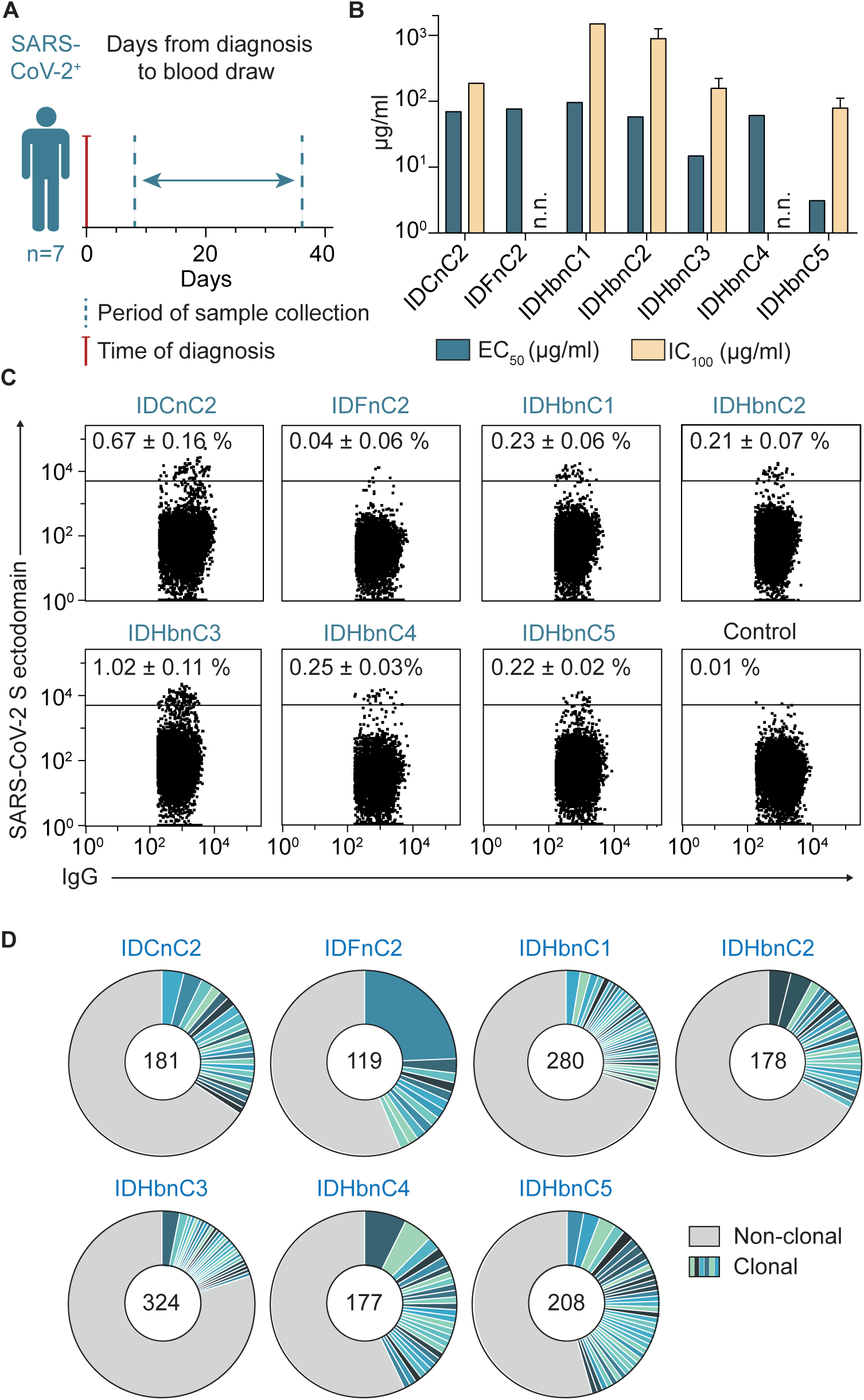
SARS-CoV-2 infection induces a polyclonal B cell and antibody response. (**A**) Scheme of cross-sectional sample collection. (**B**) Binding to the trimeric SARS-CoV-2 S-ectodomain (ELISA, EC_50_) and authentic SARS-CoV-2 neutralization activity (complete inhibition of VeroE6 cell infection, IC_100_) of cross-sectional poly-IgG samples. Bar plots show arithmetic or geometric means ± SD of duplicates or quadruplicates for EC_50_ and IC_100_, respectively. n.n., not neutralizing. (**C**) Dot plots of IgG^+^ B cell analysis. Depicted numbers (%) indicate average frequencies of S-reactive B cells across several experiments (see also **Table S2** and **Figure S1**). (**D**) Clonal relationship of S-ectodomain-reactive B cells. Individual clones are coloured in shades of blue and green. Numbers of productive heavy chain sequences are given. Clone sizes are proportional to the total number of productive heavy chains per clone.

### Longitudinal analysis of the SARS-CoV-2 antibody response

To delineate the dynamics of the SARS-CoV-2 antibody response, we obtained longitudinal blood samples from an additional five infected individuals at three time points spanning 8 to 69 days post diagnosis (**Figure 2A, Table S1**). Across the different individuals, EC_50_ (S-ectodomain binding) and IC_100_ (SARS-CoV-2 neutralization) values of plasma IgG ranged from 1.54 to 129 μg/ml and 78.8 to 1,500 μg/ml, respectively (**Figure 2B, Table S2**). For each individual, however, this response remained almost unchanged over the studied period (**Figure 2A, B**).

**Figure 2.**
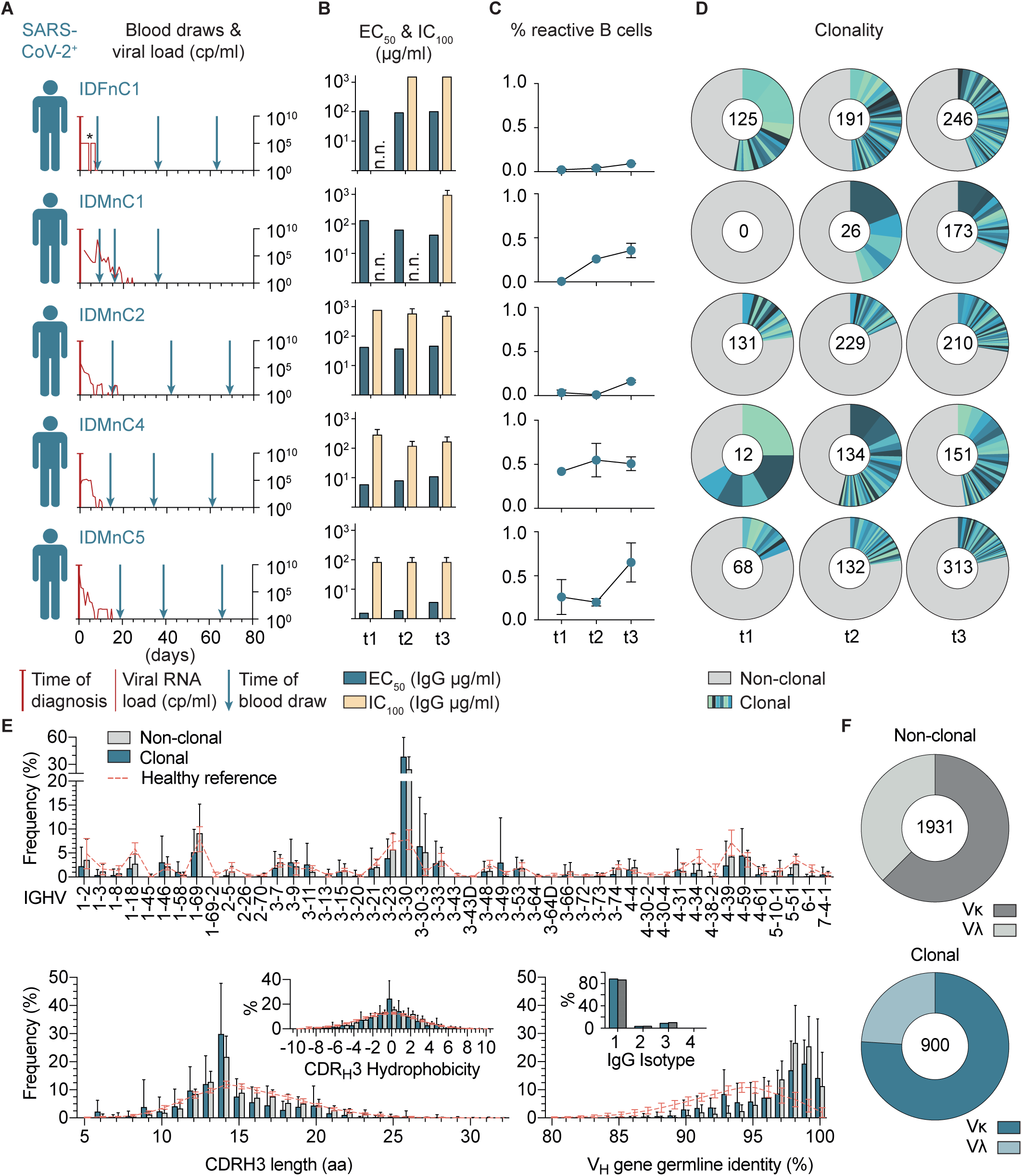
SARS-CoV-2-specific IgG^+^ B cells readily develop after infection with recurring B cell clones and a preference for the V_H_ gene segment 3-30. (**A**) Scheme of longitudinal sample collection. Viral RNA load from nasopharyngeal swabs is indicated in red (cp/ml, right Y axis). *Viral load for IDFnC1 is given as positive/negative result. (**B**) Binding to trimeric SARS-CoV-2 S-ectodomain (ELISA, EC_50_) and authentic SARS-CoV-2 neutralization activity (complete inhibition of VeroE6 cell infection, IC_100_) of longitudinal poly-IgG samples. Bar plots show arithmetic or geometric means ± SD of duplicates or quadruplicates for EC_50_ and IC_100_, respectively. n.n., not neutralizing. (**C**) Percentage of SARS-CoV-2 S-ectodomain-reactive IgG^+^ B cells over time (mean ± SD). (**D**) Clonal relationship over time. Individual clones are coloured in shades of blue and green. Numbers of productive heavy chain sequences per time point are given. (**E**) Frequencies of V_H_ gene segments (top), CDRH3 length and CDRH3 hydrophobicity (lower left), as well as V_H_ gene germline identity and IgG isotype of clonal and non-clonal sequences (lower right) from all 12 subjects and time points. NGS reference data from 48 healthy individuals (collected before the outbreak of SARS-CoV-2) are depicted in red. (**F**) Ratio of κ and λ light chains in non-clonal (top, grey) and in clonal (bottom, blue) sequences.

To investigate B cell clonality and antibody characteristics on a single cell level, we proceeded to sort S-ectodomain-reactive IgG^+^ B cells from all five subjects at the different time points (t1, t2, t3). We found up to 0.65% SARS-CoV-2-reactive B cells, with a tendency towards higher frequencies at later time points (**Figure 2C**). From a total of 2,562 B cells, we detected 254 B cell clones (**Table S3**). 51% of these clones (129) were recurrently detected, suggesting the persistence of SARS-CoV-2 reactive B cells over the investigated period of 2.5 months. When separated by individual time points, the fraction of clonally related sequences ranged from 18% to 67% across patients and remained constant or showed only moderate decreases over time (**Figure 2D**).

Next, we analysed the single cell Ig sequences (6,587 productive heavy and light chains) from all 12 patients (**Figure 2E-F, Figure S2**). Here, clonally related and non-clonal sequences similarly presented a broad spectrum of V_H_ gene segments, normally distributed heavy chain complementarity-determining region 3 (CDRH3) lengths, symmetrical CDRH3 hydrophobicity distributions, and a predominance of the IgG1 isotype (**Figure 2E**). However, in comparison to repertoire data from healthy individuals, IgV_H_ 3-30 was overrepresented and clonal sequences more often facilitated κ over λ light chains (3/4 in clonal versus 2/3 in non-clonal, p = 0.0027; **Figure 2F, Figure S2**). Finally, V_H_ genes of S-reactive B cells were on average less mutated than V_H_ genes from healthy IgG^+^ repertoires (median identity of 98.3 vs. 94.3, p < 0.0001; **Figure 2E, Figure S2**). We concluded that a SARS-CoV-2-reactive IgG^+^ B cell response readily develops after infection with the same B cell clones detectable over time and a preference for facilitating the V_H_ gene segment 3-30.

### Isolation of highly potent near-germline SARS-CoV-2-neutralizing antibodies from COVID-19 patients

To determine antibody characteristics and to isolate potent neutralizing antibodies, we cloned a total of 312 matched heavy and light chain pairs (70% clonal, 30% non-clonal) from all 12 patients. From 255 successfully produced IgG1 antibodies, 79 (31%) bound to the full trimeric S-ectodomain (Wrapp et al., 2020) with EC_50_ values ranging between 0.02 μg/ml and 5.20 μg/ml (**Figure 3A**). Of these, 30 antibodies showed SARS-CoV-2 reactivity by a commercial diagnostic system (Euroimmun IgG detection kit; **Figure 3A** and **B**, **Table S4**). Surface plasmon resonance (SPR) analyses using the RBD as analyte for 13 SARS-CoV-2 interacting antibodies gave dissociation constant (K_D_) values as low as 0.02 nM (**Table S4**). By determining the neutralization activity against authentic SARS-CoV-2, we found 28 neutralizing antibodies in 9 out of 12 patients with IC_100_ values ranging between 100 μg/ml (assay limit) and 0.04 μg/ml (**Figure 3C** and **D**). Of note, neutralizing activity was mainly detected among high affinity antibodies (**Figure 3B, Table S4**) and a positive correlation between neutralization and binding could be detected (r_s_ = 0.429, p = 0.023; **Figure 3E**).

**Figure 3.**
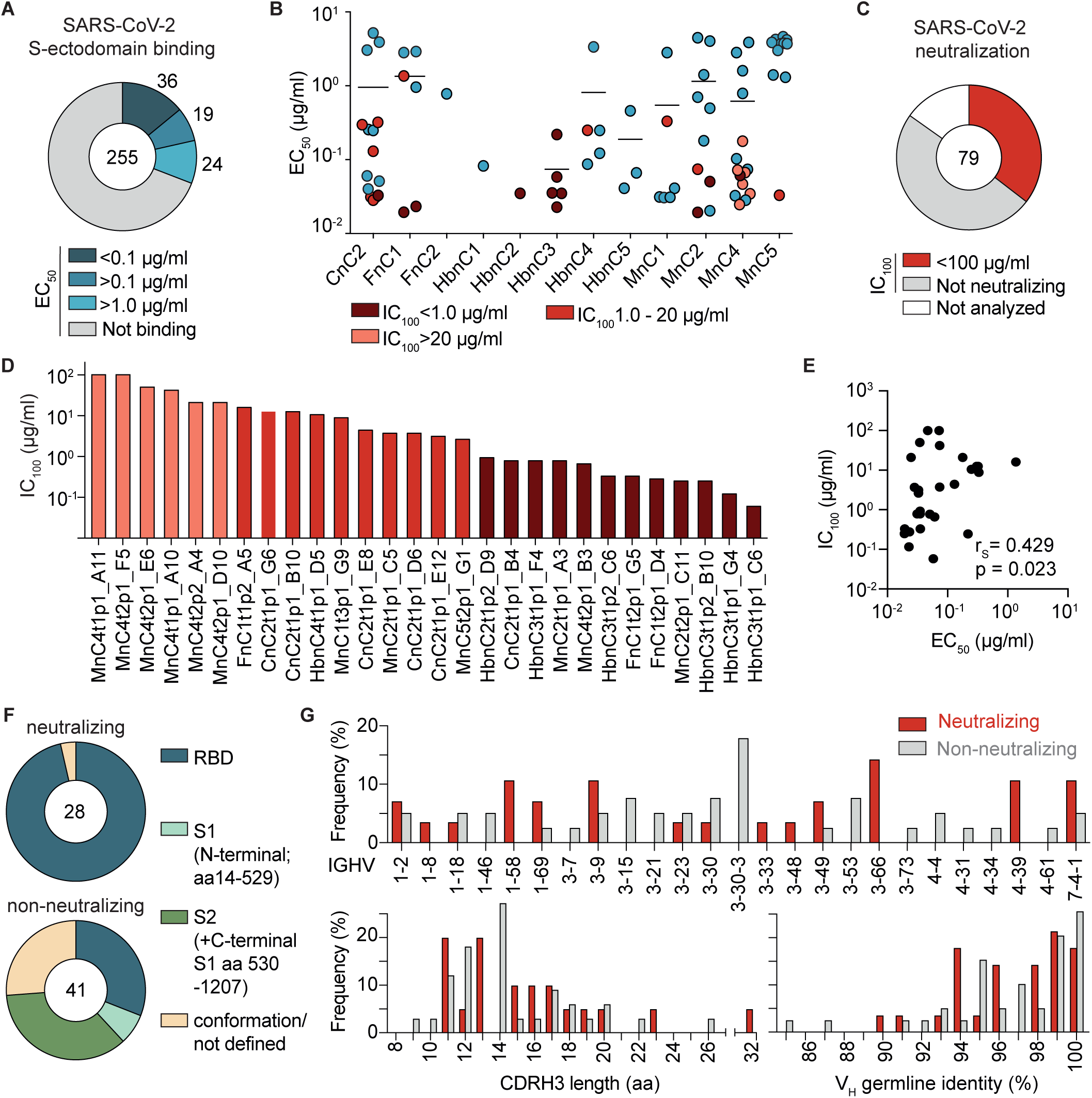
Infected individuals develop potent near-germline SARS-CoV-2-neutralizing antibodies that preferentially bind to the S-protein receptor binding domain. (**A**) Interaction of isolated antibodies with SARS-CoV-2 S-ectodomain by ELISA. Binding antibodies (blue) were defined by an EC_50_ < 30μg/ml and an OD_415-695_ > 0.25 (not shown). (**B**) EC_50_ values (mean of duplicates) of SARS-CoV-2 S-ectodomain interacting antibodies per individual. Neutralizing antibodies are labelled in shades of red. (**C**) Authentic SARS-CoV-2 neutralization activity (complete inhibition of VeroE6 cell infection, IC_100_, in quadruplicates) of S-ectodomain-specific antibodies (red). (**D**) Geometric mean potencies (IC_100_) of all neutralizing antibodies. (**E**) Correlation between S-ectodomain binding (EC_50_) and neutralization potency (IC_100_). Correlation coefficient r_S_ and approximate p-value were calculated by Spearman’s rank-order correlation. (**F**) Epitope mapping of SARS-CoV-2 S-ectodomain-specific antibodies against the RBD, truncated N-terminal S1 subunit (aa 14-529), and a monomeric S ectodomain construct by ELISA. S2 binding was defined by interaction with monomeric S but not RBD or S1. Antibodies interacting with none of the subdomains were specified as conformational epitopes or not defined. (**G**) Top: Frequencies of V_H_ gene segments for non-neutralizing and neutralizing antibodies. Clonal sequence groups were collapsed and treated as one sample for calculation of the frequencies. Bottom: CDRH3 length (left) and V_H_ gene germline identity (right) of non-neutralizing and neutralizing antibodies.

To better characterize the interaction between SARS-CoV-2 S protein and reactive antibodies, we determined binding to a truncated N-terminal S1 subunit (including the RBD), the isolated RBD, and a monomeric S ectodomain. We found 27 out of 28 neutralizing antibodies binding to the RBD, but only 31% of the non-neutralizing antibodies, suggesting that the RBD is a major site of vulnerability on the S protein. Epitopes for non-neutralizing antibodies included the N-terminal S1 domain and conformational epitopes (**Figure 3F, Table S4**). Notably, both neutralizing and non-neutralizing antibodies were characterized by a broad distribution of V_H_ as well as V_L_ gene segments and a preference for κ light chains (**Figure 3G, Figure S4**). Moreover, 32 of 79 binding and 11 of 28 neutralizing antibodies demonstrated germline identities of 99% to 100 % and no correlation was detected between neutralizing activity and the level of somatic mutation (**Figure 3G, Supplementary Table. 4, Figure S3**).

Finally, we performed a HEp-2 cell autoreactivity assay. 4 out of 28 neutralizing antibodies showed low to moderate signs of autoreactivity (**Figure S5, Table S4**) and 2 of them also reacted with other proteins (i.e. Ebola glycoprotein, HIV-1 gp140; **Table S4**). In summary, these data show that SARS-CoV-2 neutralizing antibodies develop from a broad set of different V genes and are characterized by a low degree of somatic mutations. Moreover, we were able to isolate highly potent neutralizing antibodies that present promising candidates for antibody mediated prevention and therapy of SARS-CoV-2 infection.

### Investigating ongoing somatic hypermutation in SARS-CoV-2 binding and neutralizing antibodies

To investigate the development of somatic mutations over time, we longitudinally analysed 129 recurring B cell clones that comprised 17 binding and six neutralizing antibodies. To this end, we phylogenetically matched all members of a B cell clone at a given time point with the most closely related member at the consecutive time point (331 pairings in total). Mean mutation frequencies in either direction (i.e., towards higher or lower V gene germline identities) were 0.51±0.61%, 0.08±0.51%, and 0.01±0.19% per week for all, binding, and neutralizing clonal members, respectively (**Figure 4A upper panels**). When averaging V_H_ gene germline identity of concurrent clonal members, we found a moderate increase in somatic mutations over time (**Figure 4A lower panels**). Changes were similar for binding and neutralizing subsets with one exception among the neutralizing antibodies that accumulated about 5% nucleotide mutations over the investigated period (**Figure 4A lower panel**). In line with this finding, neutralizing antibodies isolated at days 8 to 17 and days 34 to 42 post diagnosis showed V_H_ gene germline identities of 97.5% and 97.0%, respectively (**Figure 4B**). We concluded that SARS-CoV-2 neutralizing antibodies carry similar levels of somatic hypermutation independently of the time of isolation.

**Figure 4.**
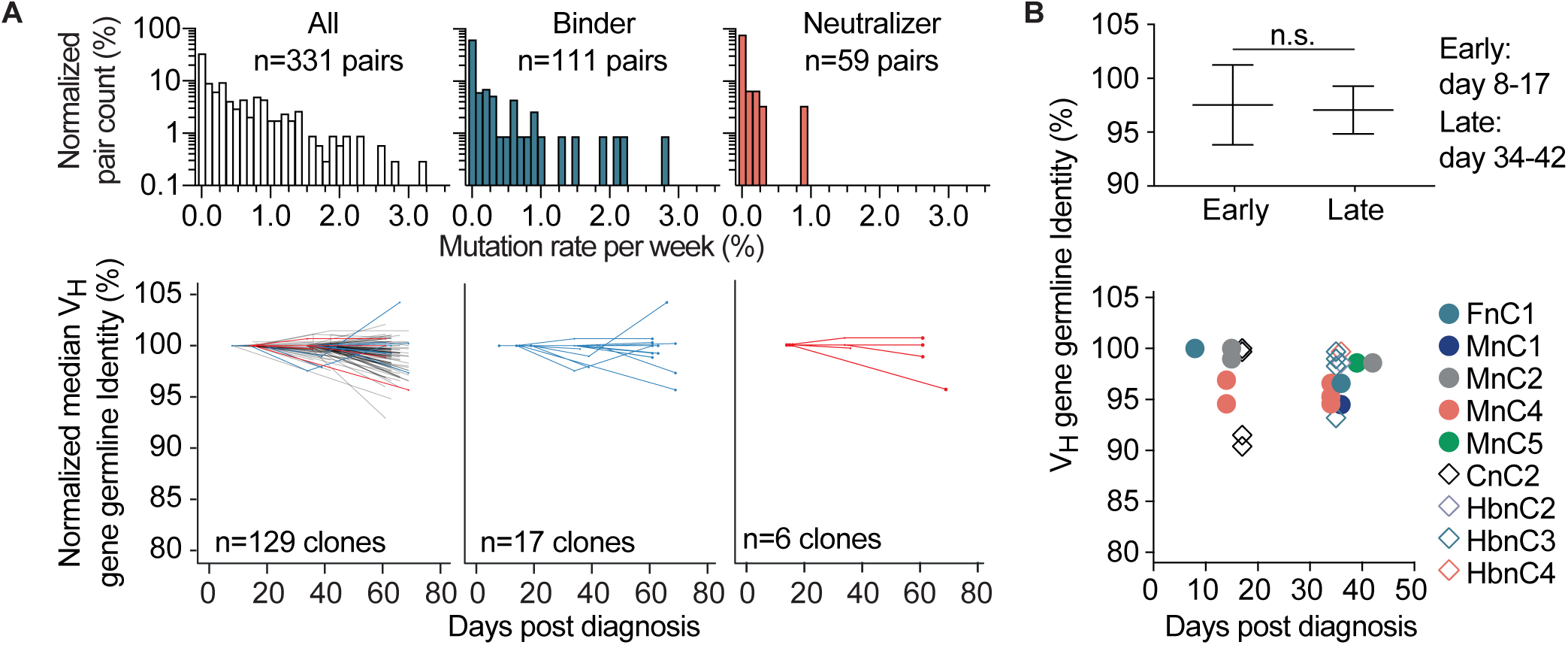
Dynamics of somatic mutations for SARS-CoV-2-specific antibodies. (**A**) Distribution of mutation rates per week for clonal members and median change in V_H_ germline identity normalized by the first measurement for each longitudinal clone. (**B**) V_H_ gene germline identity of neutralizing antibodies from different time points. Upper panel shows mean ± SD for groups of antibodies from early or late time points (two-tailed unpaired t-test). Lower panel shows V_H_ germline identities of all isolated neutralizing antibodies depending on the time between diagnosis and blood sample collection.

### Potential precursor sequences of SARS-CoV-2-neutralizing antibodies can be identified among healthy individuals

The low rate of somatic mutations in the majority of binding and neutralizing antibodies emphasizes the requirement for the presence of distinct germline recombinations in the naïve human B cell repertoire. To estimate the frequency of potential precursor B cells, we performed unbiased heavy and light chain next generation sequencing (NGS) of the naïve B cell receptor repertoires from 48 healthy donors (**Table S5**). All samples were collected before the SARS-CoV-2 outbreak and comprised a total of 1,7 million collapsed reads with 455,423 unique heavy, 170,781 κ, and 91,505 λ chain clonotypes (defined as identical V/J pairing and the same CDR3 amino acid sequence). Within this data set we searched for heavy and light chains that resemble the 79 SARS-CoV-2 binding antibodies (**Figure 5A**). For 14 out of 79 tested antibodies, we found 61 heavy chain clonotypes with identical V/J pairs and similar (± 1 aa in length and up to 3 aa differences) CDRH3s in 28 healthy individuals (**Figure 5B** and **4C**), including one exact CDRH3 match (MnC2t1p1_C12). For light chains, we identified 1,357 κ chain precursors with exact CDR3 matches that cover 41 of 62 antibodies and 109 λ chain precursors that represent 7 of 17 antibodies (**Figure 5B** and **4C**). All 48 naive repertoires included at least one κ and one λ chain precursor. When combining heavy and light chain data, we found both precursor sequences of 9 antibodies in 14 healthy individuals (**Figure 5C**). Importantly, among these potential precursor pairs, we found three potent neutralizing antibodies (CnC2t1p1_B4, HbnC3t1p1_G4, and HbnC3t1p2_B10). While the NGS repertoire data did not include pairing information of heavy and light chain combinations, we found matched heavy and light chain sequences despite small sample sizes of on average 9,500 heavy and 2,000 to 3,500 light chain clonotypes per individual. We thus conclude that potential SARS-CoV-2 binding and neutralizing antibody precursors are likely to be abundant in naïve B cell repertoires.

**Figure 5.**
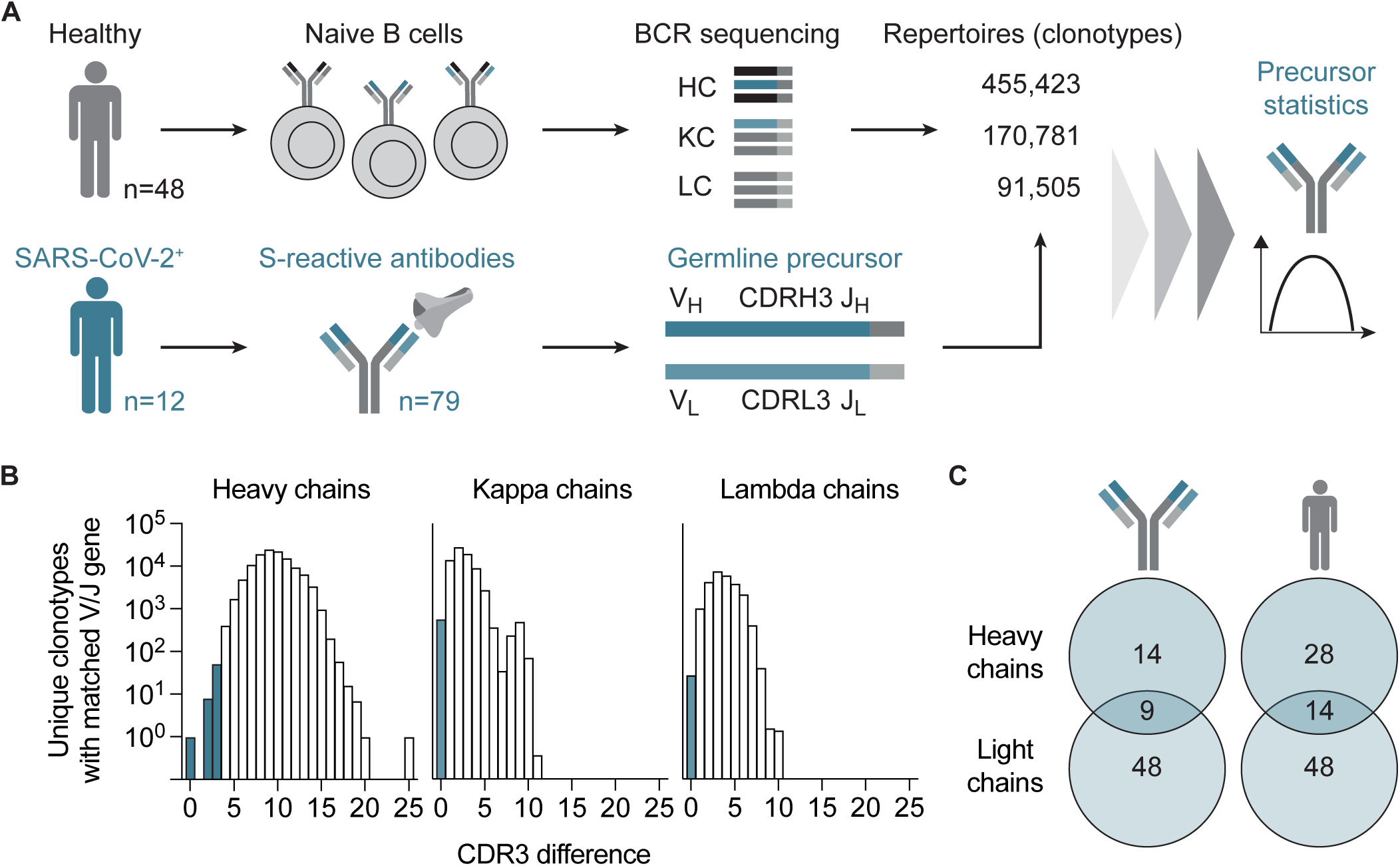
Precursor frequencies of SARS-CoV-2-specific antibodies in naïve repertoires of healthy individuals. (**A**) Strategy for precursor identification from healthy naïve B cell receptor (BCR) repertoires. HC, heavy chain; KC, κ chain; LC, λ chain; V_H_/V_L_, heavy and light chain V gene; CDRH3/CDRL3, heavy and light chain CDR3. (**B**) Number of clonotypes in healthy naïve B cell repertoires (n=48) with matched V/J genes from SARS-CoV-2 binding antibodies (n=79), plotted against the CDR3 difference. Bars of included potential precursors are highlighted in shades of blue. For heavy chains, CDR3s were allowed to differ one amino acid in length and contain up to 3 amino acid mutations. For light chains, only identical CDR3s were counted. (**C**) Number of different antibody heavy and light chains for which precursors have been identified and number of different individuals from which precursor sequences have been isolated. Numbers in overlapping circles represent matched heavy and light chain combinations.

## DISCUSSION

Neutralizing antibodies can effectively target pathogens and their induction is a key objective of vaccination strategies (Fauci and Marston, 2015; Mascola and Montefiori, 2010; Walker and Burton, 2018; Zolla-Pazner et al., 2019). A detailed understanding of the human antibody response to SARS-CoV-2 is therefore critical for the development of effective immune mediated approaches against the continuing pandemic (Burton and Walker, 2020; Koff et al., 2013; Kreer et al., 2020b). Through the single cell analysis of >4,000 SARS-CoV-2-reactive B cells from 12 infected individuals, we identified highly potent human monoclonal SARS-CoV-2-neutralizing antibodies. These antibodies block authentic viral infection at concentrations as low as 0.04 μg/ml and provide a novel option for prevention and treatment of SARS-CoV-2 infection. For many viral pathogens, the development of antibody potency is dependent on prolonged affinity maturation (Abela et al., 2019; Andrews et al., 2019; Davis et al., 2019; Wec et al., 2020). In contrast, high SARS-CoV-2-neutralizing activity can be observed for antibodies that show little if any deviation from their germline precursors outside of their CDR3s. Our longitudinal analysis of the B cell response spanning a period of more than 2.5 months after SARS-CoV-2 transmission reveals that the development of a neutralizing antibody response is followed by limited additional somatic mutation. Thus, vaccine efficacy may be more dependent on the engagement of naïve B cells rather than an extended presence of antigen to enable the accumulation of multiple antibody mutations. Importantly, we observed potential heavy and light chain precursors of potent SARS-CoV-2-neutralizing antibodies among the naïve B cell repertoires of healthy individuals that were sampled before the pandemic. Given the broad gene distribution among SARS-CoV-2-neutralizing antibodies, our findings therefore indicate the potential for a broadly active SARS-CoV-2 vaccine.

## ACKNOWLEDGEMENTS

We thank all study participants who devoted time to our research; Jan Mathis Eckert, Ralf Ortmanns, and Heidrun Schößler of the health department of Heinsberg for patient enrollment; all members of the Klein and Becker Laboratories for helpful discussion and support; Jason McLellan, Nianshuang Wang, and Daniel Wrapp for sharing the SARS-CoV-2 S-ectodomain plasmid; Florian Krammer for sharing the RBD plasmid; Simon Pöpsel and Robert Hänsel-Hertsch for helpful discussion and technical support; as well as Daniela Weiland and Nadine Henn for lab management and assistance. This work was funded by grants from the German Center for Infection Research (DZIF to F.K. and S.B.), the German Research Foundation (DFG; CRC 1279, F.K.; CRC 1310, F.K.; FOR2722, M.K.), the European Research Council (ERC-StG639961, F.K.), the German Federal Ministry of Education and Research (BMBF) within the ‘Medical Informatics Initiative’ (DIFUTURE, reference number 01ZZ1804D, S.L., N.P.), the Ben B. and Joyce E. Eisenberg Foundation (R.D.), and the Ernst I Ascher Foundation and from Natan Sharansky (R.D.).

## AUTHOR CONTRIBUTIONS

Conceptualization, F.K; Methodology, F.K., S.B., C.K., M.Z., M.S.E., L.G., C.R., S.H., S.L., N.P.; Investigation, C.K., M.Z., T.W., L.G., M.S.E., C.R., S.H., M.Kor., H.G., P.S., K.V., V.D.C., H.J., R.B., A.A., V.K., A.K., H.C.D., M.Ko., T.Wo., M.J.G.T.V., C.W.; Software, C.K., S.L., N.P.; Formal Analysis, C.K., M.Z., S.L., N.P., and F.K.; Resources, F.K., S.B., R.D.; Writing - original Draft, F.K., C.K., M.Z., T.W., H.G.; Writing - review and editing, all authors; Supervision, F.K., S.B., R.D.

## SUPPLEMENTARY FIGURE LEGENDS

**Figure S1** Gating strategy for single cell sort.

CD19^+^ B cells isolated by MACS were used and cell aggregates were excluded by FSC. Living CD20^+^ IgG^+^ cells were gated and cells with a positive SARS-CoV-2 S ectodomain staining were selected for single cell sort.

**Figure S2** Light chain characteristics of sorted single cells.

Left and middle panel: Frequencies of V_L_ gene segments of clonal and non-clonal sequences are shown (κ left, λ middle). Right panel: Ratios of κ and λ within the single sample sets in clonal and non-clonal sequences. A two-tailed ratio paired t-test was performed on κ / λ ratios to test for significance.

**Figure S3** Correlation of binding and neutralization with V_H_ gene characteristics

Correlation plots of EC_50_ values of binding or neutralizing antibodies or IC_100_ values of neutralizing antibodies with CDRH3 lengths or V_H_ germline identities. Spearman correlation coefficient r_S_ and approximate p values are given.

**Figure S4** V_L_ gene distribution in non-neutralizing and neutralizing antibodies

(**A**) Frequencies of V_L_ gene segments for non-neutralizing (left, grey) and neutralizing antibodies (right, red). Clonal sequence groups were collapsed and treated as one sample for calculation of the frequencies. (**B**) Ratio of λ and κ light chains for neutralizing (left) and non-neutralizing S-ectodomain-specific antibodies (bottom, blue).

**Figure S5** Autoreactivity of selected SARS-CoV-2 binding and neutralizing antibodies.

HEp-2 cells were incubated with SARS-CoV-2 S-ectodomain antibodies at concentrations of 100 μg/ml and analysed by indirect immunofluorescence. Representative pictures of the scoring system are shown.

## METHODS

### CONTACT FOR REAGENT AND RESOURCE SHARING

Further information and requests for resources and reagents should be directed to and will be fulfilled by the Lead Contact, Florian Klein.

### EXPERIMENTAL MODELS AND SUBJECT DETAILS

#### SARS-CoV-2 infected individuals and sample collection

Samples were obtained under a study protocol approved by the Institutional Review Board of the University of Cologne and respective local IRBs (study protocol 16-054). All participants provided written informed consent and were recruited at hospitals or as outpatients. Sites of recruitment were Munich Clinic Schwabing for IDMnC1,2,4 and 5, the University Hospital of Frankfurt for patients IDFnC1, 2, and University Hospital Cologne for patient IDCnC2. Patients IDHbnC1-5 were recruited as outpatients in the county Heinsberg.

### METHOD DETAILS

#### Isolation of peripheral blood mononuclear cells (PBMCs), plasma and total IgG from whole blood

Blood draw collection was performed using EDTA tubes and/or syringes pre-filled with heparin. PBMC isolation was performed using Leucosep centrifuge tubes (Greiner Bio-one) prefilled with density gradient separation medium (Histopaque; Sigma-Aldrich) according to the manufacturer’s instructions. Plasma was collected and stored separately. For IgG isolation, 1 ml of the collected plasma was heat-inactivated (56°C for 40 min) and incubated with Protein G Sepharose (GE Life Sciences) overnight at 4°C. The suspension was transferred to chromatography columns and washed with PBS. IgGs were eluted from Protein G using 0.1 M glycine (pH=3.0) and buffered in 0.1 M Tris (pH=8.0). For buffer exchange to PBS, 30 kDa Amicon spin membranes (Millipore) were used. Purified IgG concentration was measured using a Nanodrop (A280) and samples were stored at 4°C.

#### SARS-CoV-2 S protein expression and purification

The construct encoding the prefusion stabilized SARS-CoV-2 S ectodomain (amino acids 1−1208 of SARS-CoV-2 S; GenBank: MN908947) was kindly provided by Jason McLellan (Texas, USA) and described previously (Wrapp et al., 2020). In detail, two proline substitutions at residues 986 and 987 were introduced for prefusion state stabilization, a “GSAS” substitution at residues 682–685 to eliminate the furin cleavage site, and a C-terminal T4 fibritin trimerization motif. For purification, the protein is C-terminally fused to a TwinStrepTag and 8XHisTag. Protein production was done in HEK293-6E cells by transient transfection with polyethylenimine (PEI, Sigma-Aldrich) and 1 μg DNA per 1 mL cell culture medium at a cell density of 0.8 10^6^ cells/mL in FreeStyle 293 medium (Thermo Fisher Scientific). After 7 days of culture at 37°C and 5% CO_2_, culture supernatant was harvested and filtered using a 0.45 μm polyethersulfone (PES) filter (Thermo Fisher Scientific). Recombinant protein was purified by Strep-Tactin affinity chromatography (IBA lifescience, Göttingen Germany) according to the Strep-Tactin XT manual. Briefly, filtered medium was adjusted to pH 8 by adding 100 mL 10x Buffer W (1 M Tris/HCl, pH 8.0, 1.5 M NaCl, 10 mM EDTA, IBA lifescience) and loaded with a low pressure pump at 1 mL/min on 5 mL bedvolume Strep-Tactin resin. The column was washed with 15 column volumes (CV) 1x Buffer W (IBA lifescience) and eluted with 6 x 2.5 mL 1x Buffer BXT (IBA lifescience). Elution fractions were pooled and buffer was exchanged to PBS pH 7.4 (Thermo Fisher Scientific) by filtrating four times over 100 kDa cut-off cellulose centrifugal filter (Merck).

#### Cloning and expression of different SARS-CoV-2 S protein subunits and Ebola surface glycoprotein

The RBD of the SARS-CoV-2 spike protein (MN908947; aa:319-541) was expressed in 293T cells from a plasmid kindly provided by Florian Krammer and purified using Ni-NTA Agarose (Macherey-Nagel), as previously published (Stadlbauer et al., 2020). SARS-CoV-2 S ectodomain “monomer” without trimerization domain (MN908947; aa:1-1207) and S1 subunit (MN908947; aa:14-529) regions of the spike DNA were amplified from a synthetic gene plasmid (furin site mutated; Wrapp et al., 2020) by PCR. PCR products were cloned into a modified sleeping beauty transposon expression vector containing a C-terminal thrombin cleavage and a double Strep II purification tag. For the S1 subunit, the tag was added at the 5’ end and a BM40 signal peptide was included. For recombinant protein production, stable HEK293 EBNA cell lines were generated employing the sleeping beauty transposon system (Kowarz et al., 2015). Briefly, expression constructs were transfected into the HEK293 EBNA cells using FuGENE HD transfection reagent (Promega). After selection with puromycin, cells were induced with doxycycline. Supernatants were filtered and the recombinant proteins purified via Strep-Tactin®XT (IBA Lifescience) resin. Proteins were then eluted by biotin-containing TBS-buffer (IBA Lifescience), and dialyzed against TBS-buffer. Ebola surface glycoprotein (EBOV Makona, GenBank KJ660347) and HIV-gp140 (strain YU2), both lacking the transmembrane domain and containing a GCN4 trimerization domain, were produced and purified as previously described (Ehrhardt et al., 2019).

#### Isolation of SARS-CoV S ectodomain-specific IgG^+^ B cells

B cells were isolated from PBMCs using CD19-microbeads (Miltenyi Biotec) according to the manufacturer’s instruction. Isolated B cells were stained for 20 minutes on ice with a fluorescence staining-mix containing 4’,6-Diamidin-2-phenylindol (DAPI; Thermo Fisher Scientific), anti-human CD20-Alexa Fluor 700 (BD), anti-human IgG-APC (BD), anti-human CD27-PE (BD) and DyLight488-labeled SARS-CoV-2 spike protein (10μg/mL). Dapi^-^, CD20^+^, IgG^+^, SARC-CoV-2 spike protein positive cells were sorted using a FACSAria Fusion (Becton Dickinson) in a single cell manner into 96-well plates. All wells contained 4 μl buffer, consisting of 0.5x PBS, 0.5 U/μl RNAsin (Promega), 0.5 U/μl RNaseOUT (Thermo Fisher Scientific), and 10 mM DTT (Thermo Fisher Scientific). After sorting, plates were immediately stored at −80°C until further processing.

#### Antibody heavy/light chain amplification and sequence analysis

Single cell amplification of antibody heavy and light chains was mainly performed as previously described (Kreer et al., 2020a; Schommers et al., 2020). Briefly, reverse transcription was performed with Random Hexamers (Invitrogen), and Superscript IV (Thermo Fisher Scientific) in the presence of RNaseOUT (Thermo Fisher Sicentific) and RNasin (Promega). cDNA was used to amplify heavy and light chains using PlatinumTaq HotStart polymerase (Thermo Fisher Scientific) with 6% KB extender and optimized V gene-specific primer mixes (Kreer et al., 2020a) in a sequential semi-nested approach with minor modifications to increase throughput (Manuscript in preparation). PCR products were analyzed by gel electrophoresis for correct sizes and subjected to Sanger sequencing. For sequence analysis, chromatograms were filtered for a mean Phred score of 28 and a minimal length of 240 nucleotides (nt). Sequences were annotated with IgBLAST (Ye et al., 2013) and trimmed to extract only the variable region from FWR1 to the end of the J gene. Base calls within the variable region with a Phred score below 16 were masked and sequences with more than 15 masked nucleotides, stop codons, or frameshifts were excluded from further analyses. Clonal analysis was performed separately for each patient. All productive heavy chain sequences were grouped by identical V_H_/J_H_ gene pairs and the pairwise Levenshtein distance for their CDRH3s was determined. Starting from a random sequence, clone groups were assigned four sequences with a minimal CDRH3 amino acid identity of at least 75% (with respect to the shortest CDRH3). 100 rounds of input sequence randomization and clonal assignment were performed and the result with the lowest number of remaining unassigned (non-clonal) sequences was selected for downstream analyses. All clones were cross-validated by the investigators taking shared mutations into account. V gene usage, CDRH3 length and V gene germline identity distributions for all clonal sequences (Figure 2) were determined for all input sequences without further collapsing. CDRH3 hydrophobicity was calculated based on the Eisenberg-scale (Eisenberg et al., 1984). V gene statistics for neutralizer and non-neutralizer (Figure 3) were calculated from collapsed clonal sequences.

For longitudinal analyses on mutation frequencies of recurring clones, a multiple sequence alignment for the B cell sequences was calculated with Clustal Omega (version 1.2.3; Sievers et al., 2011) using standard parameters. From this, a phylogenetic tree of the sequences was estimated with RAxML through the raxmlGUI (version 2.0.0-beta.11; Edler et al., 2019) using the GTRGAMMA substitution model (RAxML version 8.2.12; Stamatakis, 2014). Based on the phylogenetic tree distances, all variants of a clone at a given time point were matched to variants at the consecutive time point and the slope between the pairs was computed. Hamming distances between the pairs were determined and normalized for sequence length and time difference to calculate the mean mutation frequency per day. Given the median slope per clone a one-sided Wilcoxon Signed Rank Test was applied to test whether the slopes are equal to zero, with the alternative hypothesis that the slopes are smaller than zero. For visualizing the change of V_H_ gene germline identity over time, the germline identity for each clone was normalized by its median value at the first-time measurement and the median slope was plotted.

#### Next generation sequencing and evaluation of healthy control IgG^+^ and naïve B cell repertoires

B cell receptor repertoire sequence data was generated by an unbiased template-switch-based approach as previously described (Ehrhardt et al., 2019; Schommers et al., 2020). In brief, PBMCs from 48 healthy individuals (samples taken before the SARS-CoV-2 outbreak) were enriched for CD19^+^ cells with CD19-microbeads (Miltenyi Biotec). For each individual, 100,000 CD20^+^IgG^+^ and 100,000 CD20^+^IgD^+^IgM^+^CD27^-^IgG^-^ B cells were sorted into FBS (Sigma-Aldrich) using a BD FACSAria Fusion. RNA was isolated with the RNeasy Micro Kit (Qiagen) on a QiaCube (Qiagen) instrument. cDNA was generated by template-switch reverse transcription according to the SMARTer RACE 5’/3’ manual using the SMARTScribe Reverse Transcriptase (Takara) with a template-switch oligo including an 18-nucleotide unique molecular identifier (UMI). Heavy and light chain variable regions were amplified in a constant region-specific nested PCR and amplicons were used for library preparation and Illumina MiSeq 2 x 300 bp sequencing. Raw NGS reads were pre-processed and assembled to final sequences as previously described (Ehrhardt et al., 2019). To minimize the influence of sequencing and PCR errors, NGS-derived sequences were only evaluated when UMIs were found in at least three reads. For the identification of overlapping clonotypes in healthy individuals a maximum of one amino acid length difference and three or less differences in absolute amino acid composition of CDR3s were considered as similar.

#### Cloning and production of monoclonal antibodies

Antibody cloning from 1^st^ PCR products was performed as previously described (Schommers et al., 2020) by sequence and ligation-independent cloning (SLIC; Von Boehmer et al., 2016) with a minor modification. In contrast to the published protocol, PCR amplification for SLIC assembly was performed with extended primers based on 2^nd^ PCR primers (Kreer et al., 2020a) covering the complete endogenous leader sequence of all heavy and light chain V genes (Manuscript in preparation). Variable regions with endogenous leader sequences were assembled into mammalian expression vectors for IgH, IgK, or IgL and transfected into HEK293-6E cells for expression, followed by Protein G-based purification of monoclonal antibodies from culture supernatants as previously described (Schommers et al., 2020).

#### ELISA analysis to determine antibody binding activity to SARS-CoV-2 S and subunit binding

ELISA plates (Corning 3369) were coated with 2 μg/ml of protein in PBS (SARS-CoV-2 spike ectodomain, RBD, or n-terminal truncated S1) or in 2 M Urea (SARS-CoV-2 spike ectodomain “monomer” lacking the trimerization domain) at 4°C overnight. For SARS-CoV-2 spike ectodomain ELISA, plates were blocked with 5% BSA in PBS for 60 min at RT, incubated with primary antibody in 1% BSA in PBS for 90 min, followed by anti-human IgG-HRP (Southern Biotech 2040-05) diluted 1:2500 in 1% BSA in PBS for 60 min at RT. SARS-CoV-2 spike subunit ELISAs were done following a published protocol (Stadlbauer et al., 2020). ELISAs were developed with ABTS solution (Thermo Fisher 002024) and absorbance was measured at 415 nm and 695 nm. Positive binding was defined by an OD>0.25 and an EC_50_<30 μg/ml. The commercial anti-SARS-CoV-2 ELISA kit for immunoglobulin class G was provided by Euroimmun (Euroimmun Diagnostik, Lübeck, Germany). Antibody detection was done according to manufacturer’s instructions and a concentration of 50 μg/ml of antibodies and 2 mg/ml of plasma IgG was used. The samples were tested using the automated platform Euroimmun Analyzer 1.

#### Virus neutralization test

SARS-CoV-2 neutralizing activity of poly-IgG samples or human monoclonal antibodies was investigated based on a previously published protocol for MERS-CoV^39^. Briefly, samples were serially diluted in 96-well plates starting from a concentration of 1,500 μg/ml for poly-IgG and 100 μg/ml for monoclonal antibodies. Samples were incubated for 1 h at 37°C together with 100 50% tissue culture infectious doses (TCID_50_) SARS-CoV-2 (BavPat1/2020 isolate, European Virus Archive Global # 026V-03883). Cytopathic effect (CPE) on VeroE6 cells (ATCC CRL-1586) was analysed 4 days after infection. Neutralization was defined as absence of CPE compared to virus controls. For each test, a positive control (neutralizing COVID-19 patient plasma) was used in duplicates as an inter-assay neutralization standard.

#### Surface Plasmon Resonance (SPR) measurements

For SPR measurement, the RBD was additionally purified by size exclusion chromatography (SEC) purification with a Superdex200 10/300 column (GE Healthcare). Binding of the RBD to the various mAbs was measured using single-cycle kinetics experiments with a Biacore T200 instrument (GE Healthcare). Purified mAbs were first immobilized at coupling densities of 800-1200 response units (RU) on a series S sensor chip protein A (GE Healthcare) in PBS and 0.02% sodium azide buffer. One of the four flow cells on the sensor chip was empty to serve as a blank. Soluble RBD was then injected at a series of concentrations (i.e. 0.8, 4, 20, 100, and 500 nM) in PBS at a flow rate of 60 μL/min. The sensor chip was regenerated using 10 mM Glycine-HCl pH 1.5 buffer. A 1:1 binding model was used to describe the experimental data and to derive kinetic parameters. For some mAbs, a 1:1 binding model did not provide an adequate description for binding. In these cases, we fitted a two-state binding model that assumes two binding constants due to conformational change. In these cases, we report the first binding constants (*K*_D_^1^).

#### HEp-2 Cell Assay

Monoclonal antibodies were tested at a concentration of 100 μg/ml in PBS using the NOVA Lite HEp-2 ANA Kit (Inova Diagnostics) according to the manufacturer’s instructions, including positive and negative kit controls on each substrate slide. HIV-1-reactive antibodies with known reactivity profiles were included as additional controls. Images were acquired using a DMI3000 B microscope (Leica) and an exposure time of 3.5 s, intensity of 100%, and a gain of 10.

## QUANTIFICATION AND STATISTICAL ANALYSIS

Flow cytometry analysis and quantifications were done by FlowJo10. Statistical analyses were performed using GraphPad Prism (v7), Microsoft Excel for Mac (v14.7.3), Python (v3.6.8), and R (v4.0.0).

## DATA AND SOFTWARE AVAILABILITY

All data supporting the findings of this study are available within the paper and its supplementary information files.

## REFERENCES

Abela, I.A., Kadelka, C., and Trkola, A. (2019). Correlates of broadly neutralizing antibody development. Curr. Opin. HIV AIDS.

Andrews, S.F., Chambers, M.J., Schramm, C.A., Plyler, J., Raab, J.E., Kanekiyo, M., Gillespie, R.A., Ransier, A., Darko, S., Hu, J., et al. (2019). Activation Dynamics and Immunoglobulin Evolution of Pre-existing and Newly Generated Human Memory B cell Responses to Influenza Hemagglutinin. Immunity 51, 398–410.e5.

Von Boehmer, L., Liu, C., Ackerman, S., Gitlin, A.D., Wang, Q., Gazumyan, A., and Nussenzweig, M.C. (2016). Sequencing and cloning of antigen-specific antibodies from mouse memory B cells. Nat. Protoc. 11, 1908–1923.

Burton, D.R., and Walker, L.M. (2020). Rational Vaccine Design in the Time of COVID-19. Cell Host Microbe 27, 695–698.

Burton, D.R., Walker, L.M., Phogat, S.K., Chan-Hui, P.Y., Wagner, D., Phung, P., Goss, J.L., Wrin, T., Simek, M.D., Fling, S., et al. (2009). Broad and potent neutralizing antibodies from an african donor reveal a new HIV-1 vaccine target. Science (80-.).

Corti, D., Voss, J., Gamblin, S.J., Codoni, G., Macagno, A., Jarrossay, D., Vachieri, S.G., Pinna, D., Minola, A., Vanzetta, F., et al. (2011). A neutralizing antibody selected from plasma cells that binds to group 1 and group 2 influenza A hemagglutinins. Science (80-.).

Davis, C.W., Jackson, K.J.L., McElroy, A.K., Halfmann, P., Huang, J., Chennareddy, C., Piper, A.E., Leung, Y., Albariño, C.G., Crozier, I., et al. (2019). Longitudinal Analysis of the Human B Cell Response to Ebola Virus Infection. Cell 177, 1566–1582.e17.

Dong, E., Du, H., and Gardner, L. (2020). An interactive web-based dashboard to track COVID-19 in real time. Lancet. Infect. Dis. 3099, 19–20.

Edler, D., Klein, J., Antonelli, A., and Silvestro, D. (2019). raxmlGUI 2.0 beta: a graphical interface and toolkit for phylogenetic analyses using RAxML. BioRxiv 800912.

Ehrhardt, S.A., Zehner, M., Krähling, V., Cohen-Dvashi, H., Kreer, C., Elad, N., Gruell, H., Ercanoglu, M.S., Schommers, P., Gieselmann, L., et al. (2019). Polyclonal and convergent antibody response to Ebola virus vaccine rVSV-ZEBOV. Nat. Med. 25, 1589–1600.

Eisenberg, D., Schwarz, E., Komaromy, M., and Wall, R. (1984). Analysis of membrane and surface protein sequences with the hydrophobic moment plot. J. Mol. Biol. 179, 125–142.

Fauci, A.S., and Marston, H.D. (2015). Toward an HIV vaccine: A scientific journey. Science (80-.).

Flyak, A.I., Shen, X., Murin, C.D., Turner, H.L., David, J.A., Fusco, M.L., Lampley, R., Kose, N., Ilinykh, P.A., Kuzmina, N., et al. (2016). Cross-Reactive and Potent Neutralizing Antibody Responses in Human Survivors of Natural Ebolavirus Infection. Cell.

Hoffmann, M., Kleine-Weber, H., Schroeder, S., Krüger, N., Herrler, T., Erichsen, S., Schiergens, T.S., Herrler, G., Wu, N.H., Nitsche, A., et al. (2020). SARS-CoV-2 Cell Entry Depends on ACE2 and TMPRSS2 and Is Blocked by a Clinically Proven Protease Inhibitor. Cell.

Huang, C., Wang, Y., Li, X., Ren, L., Zhao, J., Hu, Y., Zhang, L., Fan, G., Xu, J., Gu, X., et al. (2020). Clinical features of patients infected with 2019 novel coronavirus in Wuhan, China. Lancet 395, 497–506.

Huang, J., Kang, B.H., Ishida, E., Zhou, T., Griesman, T., Sheng, Z., Wu, F., Doria-Rose, N.A., Zhang, B., McKee, K., et al. (2016a). Identification of a CD4-Binding-Site Antibody to HIV that Evolved Near-Pan Neutralization Breadth. Immunity.

Huang, Y., Yu, J., Lanzi, A., Yao, X., Andrews, C.D., Tsai, L., Gajjar, M.R., Sun, M., Seaman, M.S., Padte, N.N., et al. (2016b). Engineered Bispecific Antibodies with Exquisite HIV-1-Neutralizing Activity. Cell.

Joyce, M.G., Wheatley, A.K., Thomas, P. V., Chuang, G.Y., Soto, C., Bailer, R.T., Druz, A., Georgiev, I.S., Gillespie, R.A., Kanekiyo, M., et al. (2016). Vaccine-Induced Antibodies that Neutralize Group 1 and Group 2 Influenza A Viruses. Cell.

Kallewaard, N.L., Corti, D., Collins, P.J., Neu, U., McAuliffe, J.M., Benjamin, E., Wachter-Rosati, L., Palmer-Hill, F.J., Yuan, A.Q., Walker, P.A., et al. (2016). Structure and Function Analysis of an Antibody Recognizing All Influenza A Subtypes. Cell.

Koff, W.C., Burton, D.R., Johnson, P.R., Walker, B.D., King, C.R., Nabel, G.J., Ahmed, R., Bhan, M.K., and Plotkin, S.A. (2013). Accelerating next-generation vaccine development for global disease prevention. Science (80-.). 340, 1232910–1232910.

Kowarz, E., Löscher, D., and Marschalek, R. (2015). Optimized Sleeping Beauty transposons rapidly generate stable transgenic cell lines. Biotechnol. J. 10, 647–653.

Kreer, C., Döring, M., Lehnen, N., Ercanoglu, M.S., Gieselmann, L., Luca, D., Jain, K., Schommers, P., Pfeifer, N., and Klein, F. (2020a). openPrimeR for multiplex amplification of highly diverse templates. J. Immunol. Methods 480.

Kreer, C., Gruell, H., Mora, T., Walczak, A.M., and Klein, F. (2020b). Exploiting B cell receptor analyses to inform on HIV-1 vaccination strategies. Vaccines 8.

Kwakkenbos, M.J., Diehl, S.A., Yasuda, E., Bakker, A.Q., Van Geelen, C.M.M., Lukens, M. V., Van Bleek, G.M., Widjojoatmodjo, M.N., Bogers, W.M.J.M., Mei, H., et al. (2010). Generation of stable monoclonal antibody-producing B cell receptor-positive human memory B cells by genetic programming. Nat. Med.

Mascola, J.R., and Montefiori, D.C. (2010). The Role of Antibodies in HIV Vaccines. Annu. Rev. Immunol. 28, 413–444.

Sanders, J.M., Monogue, M.L., Jodlowski, T.Z., and Cutrell, J.B. (2020). Pharmacologic Treatments for Coronavirus Disease 2019 (COVID-19): A Review. JAMA.

Saphire, E.O., Schendel, S.L., Fusco, M.L., Gangavarapu, K., Gunn, B.M., Wec, A.Z., Halfmann, P.J., Brannan, J.M., Herbert, A.S., Qiu, X., et al. (2018). Systematic Analysis of Monoclonal Antibodies against Ebola Virus GP Defines Features that Contribute to Protection. Cell.

Scheid, J.F., Mouquet, H., Ueberheide, B., Diskin, R., Klein, F., Oliveira, T.Y.K., Pietzsch, J., Fenyo, D., Abadir, A., Velinzon, K., et al. (2011). Sequence and Structural Convergence of Broad and Potent HIV Antibodies That Mimic CD4 Binding. Science (80-.).

Schommers, P., Gruell, H., Abernathy, M.E., Tran, M.K., Dingens, A.S., Gristick, H.B., Barnes, C.O., Schoofs, T., Schlotz, M., Vanshylla, K., et al. (2020). Restriction of HIV-1 Escape by a Highly Broad and Potent Neutralizing Antibody. Cell 180, 471–489.e22.

Sievers, F., Wilm, A., Dineen, D., Gibson, T.J., Karplus, K., Li, W., Lopez, R., McWilliam, H., Remmert, M., Söding, J., et al. (2011). Fast, scalable generation of high-quality protein multiple sequence alignments using Clustal Omega. Mol. Syst. Biol. 7.

Stadlbauer, D., Amanat, F., Chromikova, V., Jiang, K., Strohmeier, S., Arunkumar, G.A., Tan, J., Bhavsar, D., Capuano, C., Kirkpatrick, E., et al. (2020). SARS-CoV-2 Seroconversion in Humans: A Detailed Protocol for a Serological Assay, Antigen Production, and Test Setup. Curr. Protoc. Microbiol. 57.

Stamatakis, A. (2014). RAxML version 8:A tool for phylogenetic analysis and post-analysis of large phylogenies. Bioinformatics 30, 1312–1313.

Walker, L.M., and Burton, D.R. (2018). Passive immunotherapy of viral infections: “super-antibodies” enter the fray. Nat. Rev. Immunol. 18, 297–308.

Walls, A.C., Park, Y.J., Tortorici, M.A., Wall, A., McGuire, A.T., and Veesler, D. (2020). Structure, Function, and Antigenicity of the SARS-CoV-2 Spike Glycoprotein. Cell 181, 281–292.e6.

Wec, A.Z., Haslwanter, D., Abdiche, Y.N., Shehata, L., Pedreño-Lopez, N., Moyer, C.L., Bornholdt, Z.A., Lilov, A., Nett, J.H., Jangra, R.K., et al. (2020). Longitudinal dynamics of the human B cell response to the yellow fever 17D vaccine. Proc. Natl. Acad. Sci. U. S. A. 117, 6675–6685.

Wrapp, D., Wang, N., Corbett, K.S., Goldsmith, J.A., Hsieh, C.L., Abiona, O., Graham, B.S., and McLellan, J.S. (2020). Cryo-EM structure of the 2019-nCoV spike in the prefusion conformation. Science (80-.).

Wu, X., Yang, Z.Y., Li, Y., Hogerkorp, C.M., Schief, W.R., Seaman, M.S., Zhou, T., Schmidt, S.D., Wu, L., Xu, L., et al. (2010). Rational design of envelope identifies broadly neutralizing human monoclonal antibodies to HIV-1. Science (80-.).

Ye, J., Ma, N., Madden, T.L., and Ostell, J.M. (2013). IgBLAST: an immunoglobulin variable domain sequence analysis tool. Nucleic Acids Res.

Zhou, P., Yang, X.-L., Wang, X.-G., Hu, B., Zhang, L., Zhang, W., Si, H.-R., Zhu, Y., Li, B., Huang, C.-L., et al. (2020). A pneumonia outbreak associated with a new coronavirus of probable bat origin. Nature 579, 270–273.

Zhu, N., Zhang, D., Wang, W., Li, X., Yang, B., Song, J., Zhao, X., Huang, B., Shi, W., Lu, R., et al. (2020). A novel coronavirus from patients with pneumonia in China, 2019. N. Engl. J. Med. 382, 727–733.

Zolla-Pazner, S., Alvarez, R., Kong, X.P., and Weiss, S. (2019). Vaccine-induced V1V2-specific antibodies control and or protect against infection with HIV, SIV and SHIV. Curr. Opin. HIV AIDS.

