## Supplementary Data for "Longitudinal isolation of potent near-germline SARS-CoV-2-neutralizing antibodies from COVID-19 patients"

### Supplementary Figures


| | |
|---|---|
| Supplementary Figure 1 | Gating strategy for single cell sort. |
| Supplementary Figure 2 | Light chain characteristics of sorted single cells. |
| Supplementary Figure 3 | Correlation of binding and neutralization with VH gene characteristics |
| Supplementary Figure 4 | VL gene distribution in non-neutralizing and neutralizing antibodies |
| Supplementary Figure 5 | Autoreactivity of selected SARS-CoV-2 binding and neutralizing antibodies. |


### Supplementary Tables


| | |
|---|---|
| Supplementary Table 1 | Clinical data of SARS-CoV-2 infected study patients |
| Supplementary Table 2 | Binding and neutralization activities of poly-IgG of SARS-CoV-2 infected study patients |
| Supplementary Table 3 | B cell analysis summary |
| Supplementary Table 4 | Characteristics of isolated SARS-CoV-2 interacting antibodies |
| Supplementary Table 5 | Healthy control individuals |

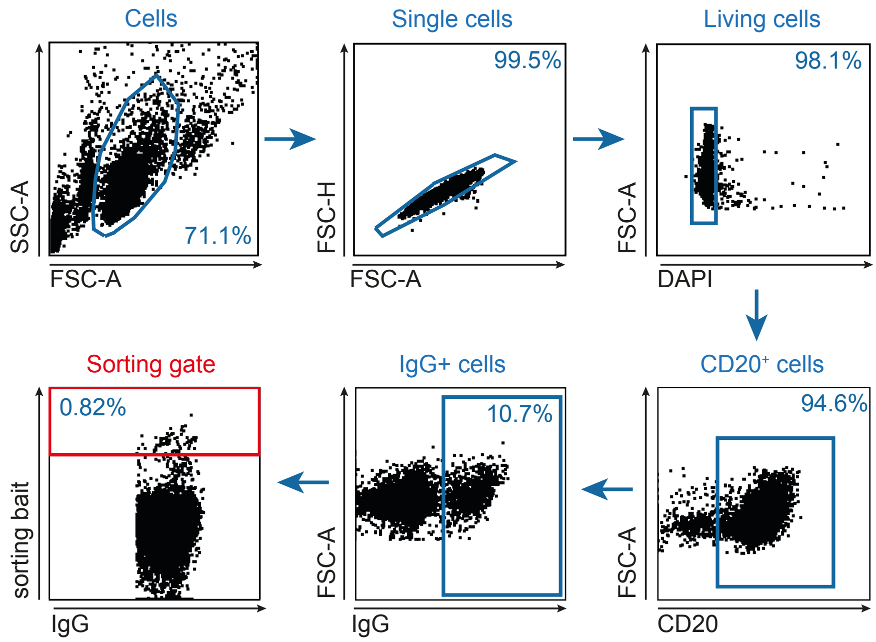


**Figure S1**. Gating strategy for single cell sort, Related to Figure 1 and Figure 2
CD19+ B cells isolated by MACS were used and cell aggregates were excluded by FSC. Living CD20+ IgG+ cells were gated and cells with a positive SARS-CoV-2 S ectodomain staining were selected for single cell sort.

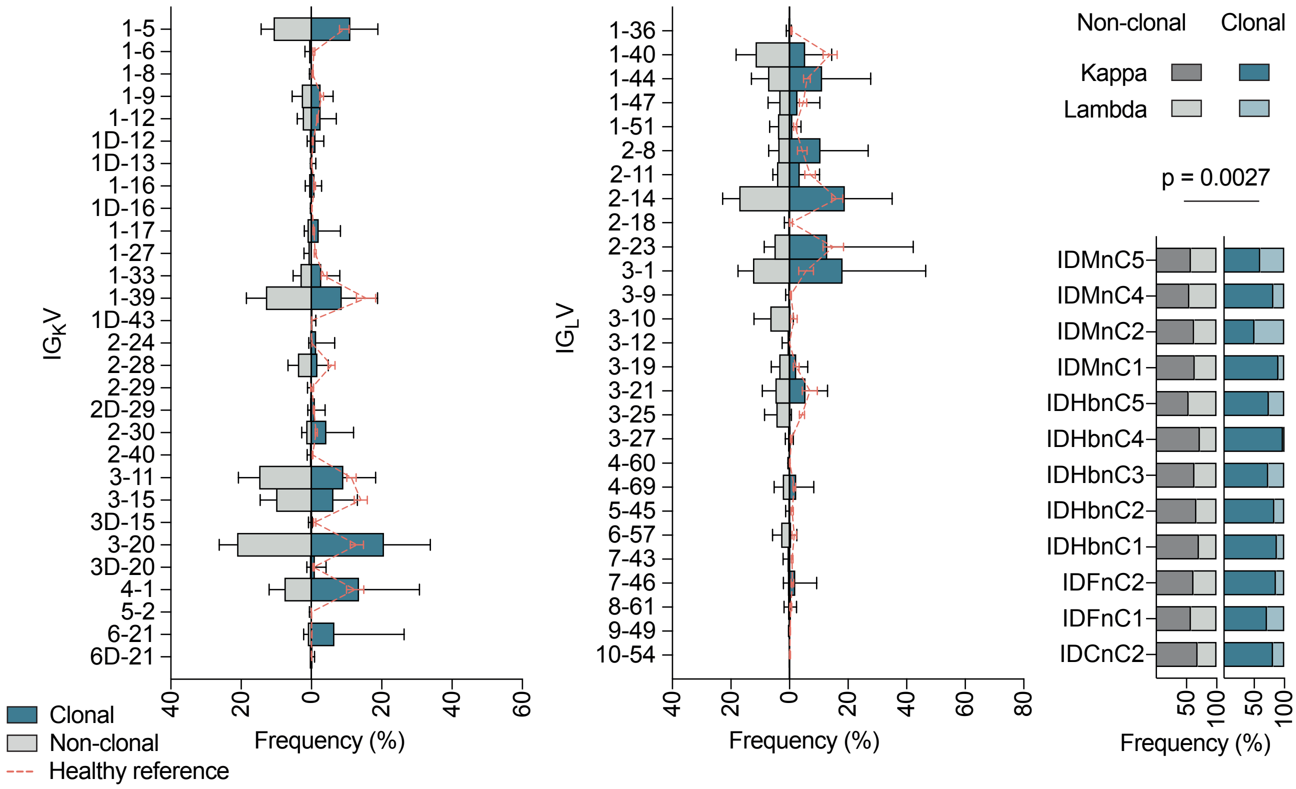


**Figure S2**. Light chain characteristics of sorted single cells, Related to Figure 2

Left and middle panel: Frequencies of VL gene segments of clonal and non-clonal sequences are shown (κ left, λ middle). Right panel: Ratios of κ and λ within the single sample sets in clonal and non-clonal sequences. A two-tailed ratio paired t-test was performed on κ / λ ratios to test for significance.

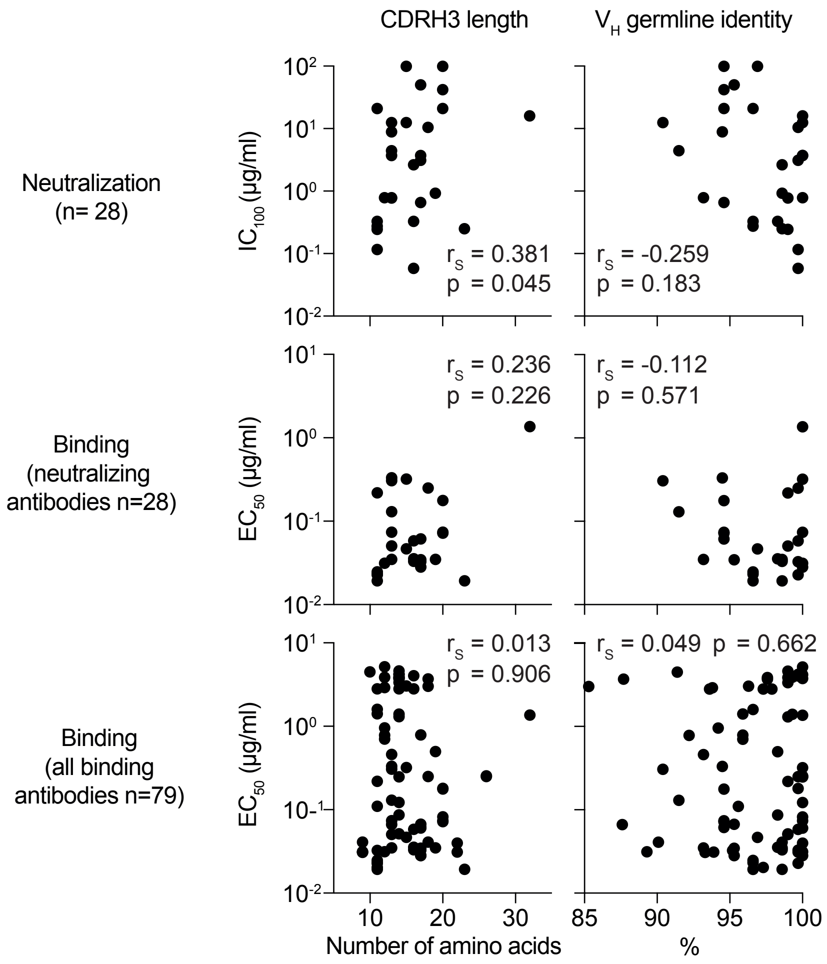


**Figure S3.** Correlation of binding and neutralization with $V_H$ gene characteristics, Related to Figure 3
Correlation plots of $EC_{50}$ values of binding or neutralizing antibodies or $IC_{100}$ values of neutralizing antibodies with CDRH3 lengths or $V_H$ germline identities. Spearman correlation coefficient $r_S$ and approximate p values are given.

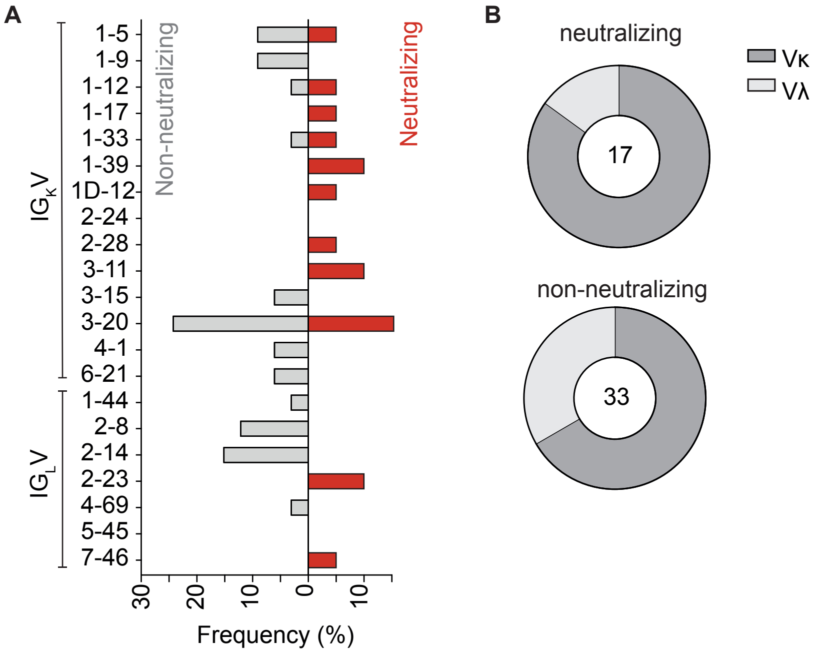


**Figure S4.** $V_L$ gene distribution in non-neutralizing and neutralizing antibodies, Related to Figure 3
(A) Frequencies of $V_L$ gene segments for non-neutralizing (left, grey) and neutralizing antibodies (right, red). Clonal sequence groups were collapsed and treated as one sample for calculation of the frequencies. (B) Ratio of λ and κ light chains for neutralizing (left) and non-neutralizing S-ectodomain-specific antibodies (bottom, blue).

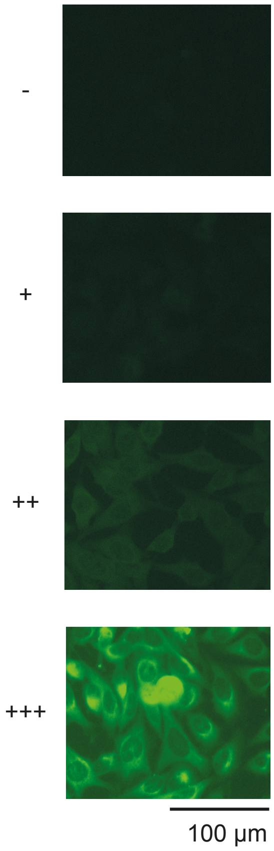


| Antibody | Result | Antibody | Result |
|---|---|---|---|
| CnC2t1p1_B4 | - | MnC1t3p1_F3 | - |
| CnC2t1p1_B10 | - | MnC1t3p1_G9 | ++ |
| CnC2t1p1_D6 | - | | |
| CnC2t1p1_E8 | - | MnC2t1p1_A3 | - |
| CnC2t1p1_E12 | - | MnC2t1p1_A12 | - |
| CnC2t1p1_F5 | ++ | MnC2t1p1_C1 | - |
| CnC2t1p1_G6 | - | MnC2t1p1_C5 | - |
| | | MnC2t1p1_D7 | - |
| FnC1t1p1_C11 | - | MnC2t2p1_C11 | - |
| FnC1t1p2_A5 | ++ | MnC2t2p1_G7 | - |
| FnC1t2p1_A12 | + | | |
| FnC1t2p1_D4 | - | MnC4t1p1_A10 | - |
| FnC1t2p1_G5 | - | MnC4t1p1_A11 | - |
| | | MnC4t2p1_B3 | - |
| HbnC2t1p2_D9 | - | MnC4t2p1_C5 | - |
| | | MnC4t2p1_D1 | - |
| HbnC3t1p1_C6 | - | MnC4t2p1_D10 | - |
| HbnC3t1p1_F4 | + | MnC4t2p1_E6 | - |
| HbnC3t1p1_G4 | ++ | MnC4t2p1_F5 | - |
| HbnC3t1p2_B10 | - | | - |
| HbnC3t1p2_C6 | - | MnC4t2p2_A4 | - |
| HbnC4t1p1_D5 | - | MnC5t1p1_C6 | - |
| | | MnC5t2p1_E9 | - |
| | | MnC5t2p1_G1 | - |
| Controls | | 3BNC117 | + |
| NIH45-46 | ++ | 2F5 | +++ |
| NIH45-46$^{G54W}$ | +++ | 4E10 | +++ |

**Figure S5.** Autoreactivity of selected SARS-CoV-2 binding and neutralizing antibodies, Related to Figure 3
HEp-2 cells were incubated with SARS-CoV-2 S-ectodomain antibodies at concentrations of 100 µg/ml and analysed by indirect immunofluorescence. Representative pictures of the scoring system are shown.

**Table S1: Clinical data of SARS-CoV-2-infected study patients**

| Patient ID | Sex | Age | Symptoms | Radiological findings | Detection of viral RNA | Days from onset of symptoms to diagnosis | Days from diganosis to blood draw 1 | Days from diganosis to blood draw 2 | Days from diganosis to blood draw 3 | Days from diagnosis to last negative PCR |
|---|---|---|---|---|---|---|---|---|---|---|
| IDCnC2 | M | 54 | dry cough, fever, arthralgia, hyposmia | N/A | NP swabs | 3 | 17 | N/A | N/A | 23 |
| IDFnC2 | M | 59 | asymptomatic | no findings | NP swabs, stool | No Symtpoms | 8 | N/A | N/A | 9 |
| IDHbnC1 | F | 52 | dry cough, fever | N/A | NP swabs | 2 | 35 | N/A | N/A | 19 |
| IDHbnC2 | F | 44 | dry cough, fever, arthralgia, hyposmia | N/A | NP swabs | 2 | 36 | N/A | N/A | 18 |
| IDHbnC3 | F | 38 | asymptomatic | N/A | NP swabs | N/A | 35 | N/A | N/A | 17 |
| IDHbnC4 | F | 50 | dry cough, fever, dyspnea | N/A | NP swabs | N/A | 36 | N/A | N/A | 16 |
| IDHbnC5 | M | 56 | dry cough, fever, dyspnea, hyposmia, ageusia, hepatitis, diarrhea | N/A | NP swabs | 2 | 36 | N/A | N/A | 26 |
| IDFnC1 | F | 45 | dry cough, rash, pharyngitis, otitis | no findings | NP swabs, stool | 13 | 8 | 36 | 63 | 8 |
| IDMnC1 (#10)* | M | 58 | dry cough, fever, hyposmia, ageusia | interstitial pneumonia of the left lower lobe | NP swabs, sputum, stool | 2 | 9 | 16 | 36 | 26 |
| IDMnC2 (#3)* | M | 28 | dry cough, fever, sinusitis, dysosmia, ageusia, athralgia | N/A | NP swabs, sputum, stool | 3 | 15 | 42 | 69 | 19 |
| IDMnC4 (#14)* | F | 49 | dry cough, fever, diarrhea | N/A | NP swabs, sputum, stool | 2 | 14 | 34 | 61 | 12 |
| IDMnC5 (#7)* | M | 52 | dry cough, fever, hyposmia, hypogeusia, dyspnea | bipulmonal, interstitial pneumonia, minimal pleural effusion | NP swabs, sputum, stool | 4 | 19 | 39 | 66 | 23 |

*Clinical data partially taken from Wolfel R (2020) Virological assessment of hospitalized patients with COVID-2019. Nature; ID in brackets corresponds to ID in this reference.
M, F : male, female
n/a : not available
NP : nasopharyngeal

**Table S2: Binding and neutralization activities of poly-IgG of SARS-CoV-2-infected study patients**

| poly-IgG | | Binding data[1] | | Neutralization |
|---|---|---|---|---|
| | | SARS-CoV-2 (spike proten) | | |
| Name | time point | EC50 (µg/ml) | Commercial ELISA* | IC100 (µg/ml) |
| IDCnC2 | 1 | 15,65 | 0,75 | 187,50 |
| IDFnC1 | 1 | 106,44 | 0,16 | n.n. |
| | 2 | 92,19 | 0,27 | 1500,00 |
| | 3 | 100,81 | 0,26 | 1500,00 |
| IDFnC2 | 1 | 76,37 | 0,17 | n.n. |
| IDHnbC1 | 1 | 96,14 | 0,27 | 1500,00 |
| IDHnbC2 | 1 | 58,05 | 0,20 | 891,91 |
| IDHnbC3 | 1 | 14,82 | 1,84 | 157,67 |
| IDHnbC4 | 1 | 61,14 | <0.25 | n.n. |
| IDHnbC5 | 1 | 3,10 | 5,75 | 78,83 |
| IDMnC1 | 1 | 129,18 | <0.25 | n.n. |
| | 2 | 61,94 | 0,16 | n.n. |
| | 3 | 41,74 | 0,33 | 891,91 |
| IDMnC2 | 1 | 41,47 | 0,43 | 750,00 |
| | 2 | 36,56 | 0,73 | 530,33 |
| | 3 | 45,60 | 0,60 | 445,95 |
| IDMnC3 | 1 | 47,48 | 0,30 | 891,91 |
| IDMnC4 | 1 | 5,97 | 3,87 | 265,17 |
| | 2 | 8,29 | 3,90 | 111,49 |
| | 3 | 10,83 | 3,03 | 157,67 |
| IDMnC5 | 1 | 1,54 | 7,75 | 78,83 |
| | 2 | 2,04 | 6,77 | 78,80 |
| | 3 | 4,10 | 5,01 | 78,83 |

[1)] Intensity above an OD of
$EC_{50}$ below 30 µg/ml was defined as binding

* Euroimmun IgG detection kit (O.D. 450nm / 620nm)

n.n. not neutralizing

**Table S3: B cell analysis summary**

| Patient ID | % IgG+ B cells† | % bait positive† | Isolated B cells | Productive Heavy chains | % clonal | Total clones | Median clone size | Largest clone size | Cloned mAbs | S-binding mAbs | neutralizing mAbs |
|---|---|---|---|---|---|---|---|---|---|---|---|
| IDCnC2 | 10.63 ± 0.12 | 0.67 ± 0.16 | 210 | 181 | 34 | 23 | 2 | 7 | 51 | 15 | 6 |
| IDFnC2 | 10.42 ± 1.1 | 0.04 ± 0.06 | 176 | 119 | 43 | 12 | 2 | 29 | 14 | 1 | 0 |
| IDHbnC1 | 6.74 ± 0.41 | 0.23 ± 0.06 | 311 | 280 | 30 | 36 | 2 | 7 | 24 | 1 | 0 |
| IDHbnC2 | 11.87 ± 0.5 | 0.21 ± 0.07 | 213 | 178 | 33 | 24 | 2 | 7 | 12 | 1 | 1 |
| IDHbnC3 | 5.98 ± 0.13 | 1.02 ± 0.11 | 372 | 324 | 20 | 27 | 2 | 10 | 15 | 5 | 5 |
| IDHbnC4 | 9.46 ± 0.22 | 0.25 ± 0.03 | 223 | 144 | 42 | 27 | 2 | 13 | 14 | 5 | 1 |
| IDHbnC5 | 15.83 ± 0.5 | 0.22 ± 0.02 | 246 | 208 | 45 | 38 | 2 | 6 | 6 | 3 | 0 |
| IDFnC1‡ | 17.1 ± 0.82 | 0.05 ± 0.03 | 652 | 562 | 47 | 95 | 2 | 23 | 39 | 6 | 3 |
| IDMnC1 (#10)*‡ | 6.06 ± 1.24 | 0.27 ± 0.16 | 245 | 199 | 34 | 20 | 2 | 22 | 19 | 6 | 1 |
| IDMnC2 (#3)*‡ | 10.75 ± 0.69 | 0.07 ± 0.07 | 732 | 570 | 22 | 49 | 2 | 13 | 30 | 10 | 3 |
| IDMnC4 (#14)*‡ | 7.12 ± 0.65 | 0.52 ± 0.15 | 348 | 297 | 51 | 40 | 2 | 18 | 19 | 15 | 7 |
| IDMnC5 (#7)*‡ | 7.32 ± 0.72 | 0.44 ± 0.28 | 585 | 513 | 21 | 50 | 2 | 5 | 12 | 11 | 1 |
| **mean** | 9.94 ± 3.66 | 0.33 ± 0.29 | 359 | 298 | 35 | 37 | 2 | 13 | 21 | 7 | 2 |
| **sum** | - | - | 4313 | 3575 | - | 441 | - | - | 255 | 79 | 28 |

* ID in brackets corresponds to IDs in Wolfel R (2020) Virological assessment of hospitalized patients with COVID-2019. Nature.
† %IgG- and bait positive cells show mean ± std over all sorted plates
‡ For longitudinal data % IgG and % bait positives are given as mean values and clonal analysis was performed over all time points. See to Figure 2 for time-resolved analyses.

**Table S4: Characteristics of isolated SARS-CoV-2-interacting antibodies**

| Antibody | | | | | | | Binding data[1] | | | | Epitope | | | HEp2 autoreactivity | Neutralization |
|---|---|---|---|---|---|---|---|---|---|---|---|---|---|---|---|
| Name | Clone # | Number of clonal members | VH-gene | VH-gene germline identity | VL-gene | VL-gene germline identity | SARS-CoV-2 (spike proten) $EC_{50}$ (µg/ml) | SARS-CoV-2 (spike proten) $K_D$ (nM) | SARS-CoV-2 (spike proten) Commercial ELISA* | EBOVΔTM $EC_{50}$ | RBD[2] $EC_{50}$ | S1[3] $EC_{50}$ | monomeric $EC_{50}$ | | IC100 (µg/ml) |
| CnC2t1p1_A2 | C2_5 | 4 | IGHV1-46 | 99,7 | IGKV3-20 | 100 | 0,03 | N/A | 2,40 | - | + | ++ | ++ | N/A | n.n. |
| CnC2t1p1_A6 | C2_17 | 2 | IGHV3-30 | 99,3 | IGKV3-11 | 100 | 3,90 | N/A | N/A | - | - | + | ++ | N/A | N/A |
| CnC2t1p1_B10 | C2_8 | 2 | IGHV1-69 | 100 | IGKV3-11 | 100 | 0,32 | N/A | N/A | - | +++ | +++ | +++ | - | 12,50 |
| CnC2t1p1_B4 | C2_2 | 2 | IGHV1-18 | 100 | IGLV2-23 | 100 | 0,03 | N/A | 7,57 | - | +++ | +++ | +++ | - | 0,78 |
| CnC2t1p1_D6 | C2_23 | 2 | IGHV3-49 | 100 | IGKV2-28 | 98 | 0,03 | N/A | 9,26 | - | +++ | +++ | +++ | - | 3,72 |
| CnC2t1p1_E11 | single | - | IGHV3-30 | 100 | IGLV2-14 | 99,3 | 5,20 | N/A | <0.15 | - | - | + | + | N/A | n.n. |
| CnC2t1p1_E12 | C2_23 | 2 | IGHV3-49 | 99,7 | IGKV2-28 | 96 | 0,03 | N/A | 9,03 | - | +++ | +++ | +++ | - | 3,13 |
| CnC2t1p1_E7 | C2_5 | 4 | IGHV1-46 | 100 | IGKV3-20 | 99 | 0,04 | N/A | N/A | - | - | ++ | + | N/A | n.n. |
| CnC2t1p1_E8 | C2_3 | 6 | IGHV1-2 | 91,5 | IGLV2-23 | 95,6 | 0,13 | N/A | 6,88 | - | +++ | +++ | +++ | - | 4,42 |
| CnC2t1p1_F5 | single | - | IGHV3-30 | 99 | IGLV5-45 | 99,7 | 0,05 | N/A | N/A | +++ | + | + | +++ | ++ | N/A |
| CnC2t1p1_G6 | C2_3 | 6 | IGHV1-2 | 90,4 | IGLV2-23 | 94,6 | 0,31 | N/A | 4,72 | - | +++ | +++ | +++ | - | 12,50 |
| CnC2t1p2_E4 | C2_11 | 3 | IGHV3-15 | 100 | IGKV3-20 | 100 | 0,26 | N/A | <0.15 | - | - | + | + | N/A | n.n. |
| CnC2t1p2_F12 | C2_10 | 2 | IGHV3-15 | 100 | IGKV3-20 | 99,3 | 0,06 | N/A | <0.15 | - | - | + | + | N/A | n.n. |
| CnC2t1p2_F4 | C2_15 | 3 | IGHV3-30 | 96,3 | IGKV6-21 | 93,4 | 3,04 | N/A | <0.15 | - | - | + | + | N/A | n.n. |
| CnC2t1p2_G4 | C2_11 | 3 | IGHV3-15 | 100 | IGKV3-20 | 100 | 0,25 | N/A | <0.15 | - | - | + | + | N/A | n.n. |
| FnC1t1p1_B6 | F1_77 | 11 | IGHV3-53 | 94,2 | IGLV2-8 | 98,6 | 0,96 | N/A | <0.15 | - | +++ | ++ | + | N/A | n.n. |
| FnC1t1p1_C11 | F1_76 | 4 | IGHV3-53 | 93,8 | IGLV2-8 | 99,7 | 2,91 | N/A | <0.15 | + | + | + | + | - | n.n. |
| FnC1t1p2_A5 | single | - | IGHV1-8 | 100 | IGKV3-20 | 100 | 1,36 | N/A | <0.15 | + | - | + | + | ++ | 16,00 |
| FnC1t2p1_A12 | F1_89 | 2 | IGHV4-4 | 97,9 | IGLV2-14 | 97,6 | 2,81 | N/A | <0.15 | + | - | + | + | + | n.n. |
| FnC1t2p1_D4 | F1_95 | 2 | IGHV7-4-1 | 96,6 | IGKV1-33 | 99 | 0,02 | 0,20 | 9,00 | - | +++ | +++ | +++ | - | 0,28 |
| FnC1t2p1_G5 | F1_95 | 2 | IGHV7-4-1 | 96,6 | IGKV1-33 | 99 | 0,02 | 0,10 | 9,00 | - | +++ | +++ | +++ | - | 0,33 |
| FnC2t1p1_B1 | F2_12 | 3 | IGHV4-34 | 92,2 | IGLV1-44 | 92,7 | 0,78 | N/A | <0.15 | - | ++ | ++ | + | N/A | n.n. |
| HbnC1t1p3_H8 | H1_6 | 2 | IGHV3-21 | 100 | IGKV1-5 | 100 | 0,08 | N/A | <0.15 | - | - | + | +++ | N/A | n.n. |
| HbnC2t1p2_D9 | H2_20 | 2 | IGHV3-33 | 98,6 | IGKV3-11 | 99 | 0,04 | N/A | N/A | - | ++ | ++ | ++ | - | 0,93 |
| HbnC3t1p1_C6 | H3_3 | 2 | IGHV1-58 | 99,7 | IGKV3-20 | 99,7 | 0,06 | N/A | 9,00 | - | +++ | +++ | +++ | - | 0,04 |
| HbnC3t1p1_F4 | H3_10 | 10 | IGHV3-30 | 93,2 | IGKV1-5 | 96,1 | 0,04 | N/A | 8,78 | - | ++ | +++ | ++ | + | 0,78 |
| HbnC3t1p1_G4 | single | - | IGHV3-66 | 99,7 | IGKV3-20 | 98,3 | 0,02 | N/A | N/A | - | +++ | +++ | +++ | ++ | 0,12 |
| HbnC3t1p2_B10 | single | - | IGHV3-66 | 99 | IGKV3-20 | 97,9 | 0,22 | N/A | 9,00 | - | +++ | +++ | +++ | - | 0,25 |
| HbnC3t1p2_C6 | H3_3 | 2 | IGHV1-58 | 98,3 | IGKV3-20 | 99,7 | 0,04 | N/A | 10,00 | - | +++ | +++ | +++ | - | 0,33 |
| HbnC4t1p1_A1 | H4_5 | 9 | IGHV3-23 | 100 | IGKV1-33 | 100 | 0,12 | N/A | 2,98 | - | +++ | +++ | +++ | N/A | n.n. |
| HbnC4t1p1_A6 | H4_23 | 2 | IGHV3-30-3 | 99 | IGKV4-1 | 98,4 | 3,36 | N/A | <0.15 | - | - | + | + | N/A | N/A |
| HbnC4t1p1_C11 | H4_5 | 9 | IGHV3-23 | 100 | IGKV1-33 | 100 | 0,25 | N/A | 8,04 | - | +++ | +++ | +++ | N/A | n.n. |
| HbnC4t1p1_D5 | H4_27 | 4 | IGHV3-9 | 99,7 | IGKV1-39 | 98,6 | 0,25 | N/A | <0.15 | ++ | ++ | ++ | + | - | 10,51 |
| HbnC4t1p1_G2 | single | - | IGHV3-30 | 98,3 | IGKV1-39 | 99,7 | 0,09 | N/A | <0.15 | - | - | + | + | N/A | N/A |
| HbnC5t1p1_B6 | H5_2 | 6 | IGHV1-46 | 87,6 | IGKV2-24 | 97 | 0,07 | N/A | 9,86 | - | - | +++ | + | N/A | N/A |
| HbnC5t1p1_C11 | H5_4 | 2 | IGHV1-46 | 93,2 | IGKV2-24 | 96,4 | 0,46 | N/A | N/A | - | - | + | + | N/A | N/A |
| HbnC5t1p1_D4 | H5_26 | 2 | IGHV3-53 | 98,6 | IGKV1-9 | 100 | 0,04 | N/A | <0.15 | - | - | + | + | N/A | n.n. |
| MnC1t2p1_A5 | M1_19 | 22 | IGHV3-49 | 89,3 | IGKV4-1 | 96,4 | 0,03 | N/A | <0.15 | - | - | + | +++ | N/A | N/A |
| MnC1t2p1_B11 | M1_19 | 22 | IGHV3-49 | 93,3 | IGKV4-1 | 97,7 | 0,03 | N/A | <0.15 | - | - | + | +++ | N/A | n.n. |
| MnC1t3p1_B7 | M1_19 | 22 | IGHV3-49 | 90,1 | IGKV4-1 | 95,1 | 0,04 | N/A | N/A | N/A | N/A | N/A | N/A | N/A | N/A |

1) Intensity above an OD of >/=0.25 (not shown) and $EC_{50}$ below 30 µg/ml was defined as binding

2) Receptor binding domain

3) truncated S1 subunit

* Euroimmun IgG detection kit

+ $EC_{50}$ >1 µg/µl

++ 0.1µg/µl < $EC_{50}$ <1 µg/µl

+++ $EC_{50}$ <0.1 µg/µl

n.n. not neutralizing

### Table S4 (continued): Characteristics of isolated SARS-CoV-2-interacting antibodies

| Antibody | | | | | | | Binding data | | | | Epitope | | | HEp2 autoreactivity | Neutralization |
|---|---|---|---|---|---|---|---|---|---|---|---|---|---|---|---|
| Name | Clone # | Number of clonal members | VH-gene | VH-gene germline identity | VL-gene | VL-gene germline identity | SARS-CoV-2 (spike proten) EC$_{50}$ (µg/ml) | SARS-CoV-2 (spike proten) K$_D$ (nM) | SARS-CoV-2 (spike proten) IgG detection test* | EBOVΔTM EC$_{50}$ | RBD[2] EC$_{50}$ | S1[3] EC$_{50}$ | monomeric EC$_{50}$ | | IC100 (µg/ml) |
| MnC1t3p1_E3 | M1_19 | 22 | IGHV3-49 | 93,9 | IGKV4-1 | 98,7 | 0,03 | N/A | N/A | - | - | - | - | N/A | N/A |
| MnC1t3p1_F3 | single | - | IGHV3-30 | 97,3 | IGLV2-23 | 98,3 | 2,82 | N/A | N/A | + | - | - | - | - | N/A |
| MnC1t3p1_G9 | M1_2 | 3 | IGHV3-23 | 94,5 | IGLV7-46 | 97,6 | 0,33 | N/A | 5,42 | - | +++ | +++ | +++ | ++ | 8,84 |
| MnC2t1p1_A12 | single | - | IGHV1-2 | 99,7 | IGLV2-14 | 99 | 4,03 | N/A | <0.15 | - | - | - | + | - | n.n. |
| MnC2t1p1_A3 | M2_25 | 3 | IGHV3-66 | 99 | IGKV1D-12 | 100 | 0,05 | 0,70 | 6,44 | - | +++ | +++ | +++ | - | 0,78 |
| MnC2t1p1_C1 | single | - | IGHV3-21 | 99,7 | IGKV3-15 | 99 | 0,18 | N/A | <0.15 | ++ | - | ++ | +++ | - | n.n. |
| MnC2t1p1_C12 | single | - | IGHV4-31 | 97,3 | IGKV3-20 | 96,8 | 0,02 | N/A | <0.15 | - | - | - | +++ | N/A | n.n. |
| MnC2t1p1_C5 | M2_25 | 3 | IGHV3-66 | 100 | IGKV1D-12 | 100 | 0,07 | N/A | 5,85 | - | +++ | +++ | +++ | - | 3,72 |
| MnC2t2p1_C11 | single | - | IGHV1-69 | 98,6 | IGKV1-39 | 98,9 | 0,02 | 0,02 | 6,79 | - | +++ | +++ | +++ | - | 0,25 |
| MnC2t2p1_D7 | M2_29 | 3 | IGHV3-7 | 95,9 | IGLV2-8 | 98,3 | 1,41 | N/A | <0.15 | + | - | - | - | N/A | n.n. |
| MnC2t2p1_E1 | single | - | IGHV4-61 | 98,3 | IGKV1-9 | 98,6 | 0,50 | N/A | <0.15 | - | - | - | + | N/A | n.n. |
| MnC2t2p1_E4 | M2_42 | 3 | IGHV4-4 | 95,9 | IGLV2-14 | 95,9 | 0,70 | N/A | <0.15 | - | - | - | ++ | N/A | n.n. |
| MnC2t2p1_G7 | M2_33 | 2 | IGHV3-73 | 91,4 | IGLV2-14 | 93,8 | 4,48 | N/A | <0.15 | + | - | - | - | - | n.n. |
| MnC4t1p1_A10 | M4_27 | 18 | IGHV4-39 | 94,6 | IGKV1-17 | 96,8 | 0,07 | N/A | 4,87 | - | +++ | +++ | +++ | - | 42,04 |
| MnC4t1p1_A11 | M4_22 | 5 | IGHV3-48 | 96,9 | IGKV3-20 | 97,9 | 0,05 | 2000 | 4,52 | - | +++ | +++ | +++ | - | 100,00 |
| MnC4t1p1_A6 | M4_25 | 15 | IGHV3-9 | 95,3 | IGKV1-12 | 97,5 | 0,07 | N/A | 10,24 | - | +++ | +++ | +++ | N/A | n.n. |
| MnC4t1p1_A7 | M4_38 | 7 | IGHV4-61 | 93,6 | IGKV1-39 | 92,6 | 2,81 | N/A | <0.15 | - | - | - | - | N/A | N/A |
| MnC4t2p1_A8 | M4_40 | 3 | IGHV7-4-1 | 96,6 | IGKV3-20 | 96,8 | 1,60 | N/A | 0,24 | - | +++ | ++ | ++ | N/A | n.n. |
| MnC4t2p1_B3 | M4_25 | 15 | IGHV3-9 | 94,6 | IGKV1-12 | 96,8 | 0,06 | 0,09 | N/A | - | +++ | +++ | +++ | - | 0,66 |
| MnC4t2p1_C5 | M4_3 | 7 | IGHV1-2 | 95,9 | IGLV2-8 | 98,6 | 0,79 | N/A | <0.15 | ++ | + | + | + | - | n.n. |
| MnC4t2p1_D1 | single | - | IGHV1-69 | 99 | IGKV1-5 | 98,9 | 3,86 | N/A | <0.15 | + | - | - | + | - | n.n. |
| MnC4t2p1_D10 | M4_27 | 18 | IGHV4-39 | 94,6 | IGKV1-17 | 96,8 | 0,18 | 440 | 5,30 | - | +++ | +++ | +++ | - | 21,02 |
| MnC4t2p1_E6 | M4_25 | 15 | IGHV3-9 | 95,3 | IGKV1-12 | 97,5 | 0,03 | N/A | 8,08 | - | +++ | +++ | +++ | - | 50,00 |
| MnC4t2p1_F5 | M4_27 | 18 | IGHV4-39 | 94,6 | IGKV1-17 | 96,8 | 0,07 | N/A | 5,19 | - | +++ | +++ | +++ | - | 100,00 |
| MnC4t2p1_F8 | M4_25 | 15 | IGHV3-9 | 95,3 | IGKV1-12 | 97,5 | 0,03 | N/A | 10,08 | - | +++ | +++ | +++ | N/A | n.n. |
| MnC4t2p1_G3 | M4_40 | 3 | IGHV7-4-1 | 95,2 | IGKV3-20 | 95,8 | 0,03 | N/A | 7,60 | - | +++ | +++ | +++ | N/A | n.n. |
| MnC4t2p2_A4 | M4_39 | 5 | IGHV7-4-1 | 96,6 | IGKV3-20 | 95,8 | 0,02 | 14 | 10,17 | - | +++ | +++ | +++ | - | 21,02 |
| MnC4t2p2_A7 | M4_35 | 2 | IGHV4-59 | 95,6 | IGKV4-1 | 98 | 0,11 | N/A | <0.15 | - | - | - | - | N/A | N/A |
| MnC5t1p1_C12 | single | - | IGHV3-30 | 99,3 | IGKV1-9 | 99,7 | 1,40 | N/A | <0.15 | - | - | - | +++ | N/A | n.n. |
| MnC5t1p1_C4 | M5_30 | 2 | IGHV3-30-3 | 99 | IGKV3-20 | 99,3 | 1,31 | N/A | <0.15 | - | - | - | +++ | N/A | n.n. |
| MnC5t1p1_C6 | single | - | IGHV3-30-3 | 99,7 | IGKV3-20 | 100 | 4,23 | N/A | <0.15 | + | - | - | ++ | - | n.n. |
| MnC5t1p1_C7 | single | - | IGHV3-30-3 | 99 | IGKV1-5 | 99,6 | 4,62 | N/A | <0.15 | - | - | - | - | N/A | n.n. |
| MnC5t2p1_B5 | single | - | IGHV3-30-3 | 97,6 | IGKV3-20 | 99,3 | 3,69 | N/A | <0.15 | - | - | - | +++ | N/A | n.n. |
| MnC5t2p1_C2 | single | - | IGHV3-30-3 | 100 | IGLV4-69 | 100 | 4,17 | N/A | <0.15 | - | - | - | - | N/A | n.n. |
| MnC5t2p1_C5 | single | - | IGHV3-30-3 | 100 | IGKV4-1 | 100 | 3,76 | N/A | <0.15 | - | - | - | - | N/A | n.n. |
| MnC5t2p1_E12 | single | - | IGHV3-30-3 | 97,6 | IGKV6-21 | 99,3 | 3,87 | N/A | <0.15 | - | - | - | +++ | N/A | n.n. |
| MnC5t2p1_E9 | M5_1 | 4 | IGHV1-18 | 85,3 | IGKV3-15 | 97,6 | 3,01 | N/A | <0.15 | + | - | - | +++ | - | n.n. |
| MnC5t2p1_G1 | M5_4 | 2 | IGHV1-58 | 98,6 | IGKV3-20 | 99 | 0,03 | 17 | 10,48 | - | +++ | +++ | +++ | - | 2,63 |
| MnC5t2p1_G6 | M5_1 | 4 | IGHV1-18 | 87,7 | IGKV3-15 | 96,2 | 3,71 | N/A | <0.15 | - | - | - | +++ | N/A | n.n. |

1) Intensity above an OD of >/=0.25 (not shown) and EC$_{50}$ below 30 µg/ml was defined as binding

2) Receptor binding domain

3) truncated S1 subunit

* Euroimmun IgG detection kit

+ EC$_{50}$ >1 µg/µl

++ 0.1µg/µl < EC$_{50}$ <1 µg/µl

+++ EC$_{50}$ <0.1 µg/µl

n.n. not neutralizing

**Table S5: Healthy control individuals**

| Patient ID | Sex | Age | Date of blood draw |
|---|---|---|---|
| 75 | F | 47 | 27.12.16 |
| 76 | F | 51 | 27.12.16 |
| 84 | M | 47 | 28.12.16 |
| 85 | M | 23 | 28.12.16 |
| 86 | M | 47 | 28.12.16 |
| 90 | F | 25 | 02.01.17 |
| 91 | M | 55 | 02.01.17 |
| 92 | F | 54 | 02.01.17 |
| 93 | M | 34 | 02.01.17 |
| 94 | M | 31 | 02.01.17 |
| 95 | M | 19 | 03.01.17 |
| 96 | M | 55 | 03.01.17 |
| 97 | F | 22 | 03.01.17 |
| 98 | M | 34 | 03.01.17 |
| 99 | M | 51 | 03.01.17 |
| 100 | M | 46 | 03.01.17 |
| 101 | M | 46 | 04.01.17 |
| 102 | M | 31 | 04.01.17 |
| 103 | M | 25 | 04.01.17 |
| 104 | M | 21 | 04.01.17 |
| 105 | M | 20 | 04.01.17 |
| 106 | F | 48 | 04.01.17 |
| 117 | F | 24 | 10.01.17 |
| 118 | M | 30 | 10.01.17 |
| 119 | F | 28 | 10.01.17 |
| 120 | M | 21 | 10.01.17 |
| 121 | F | 21 | 10.01.17 |
| 126 | F | 24 | 11.01.17 |
| 127 | M | 42 | 11.01.17 |
| 128 | F | 67 | 11.01.17 |
| 130 | M | 37 | 12.01.17 |
| 131 | F | 67 | 12.01.17 |
| 132 | M | 39 | 12.01.17 |
| 133 | M | 35 | 12.01.17 |
| 134 | M | 36 | 12.01.17 |
| 135 | F | 22 | 18.01.17 |
| 136 | F | 20 | 18.01.17 |
| 137 | F | 21 | 18.01.17 |
| 138 | F | 26 | 18.01.17 |
| 139 | F | 36 | 18.01.17 |
| 140 | M | 23 | 18.01.17 |
| 141 | M | 21 | 24.01.17 |
| 145 | F | 21 | 24.01.17 |
| 146 | F | 21 | 24.01.17 |
| 147 | F | 50 | 01.02.17 |
| 148 | F | 24 | 01.02.17 |
| 151 | F | 49 | 14.02.17 |
| 176 | F | 20 | 09.10.17 |

M, F : male, female
